# Discovery and genomics of H_2_-oxidizing/O_2_-reducing *Deferribacterota* ectosymbiotic with protists in the guts of termites and a *Cryptocercus* cockroach

**DOI:** 10.1101/2025.08.05.668610

**Authors:** Naoya Maruoka, Rinpei Kudo, Katsura Igai, Michiru Shimizu, Masahiro Yuki, Moriya Ohkuma, Yuichi Hongoh

**Affiliations:** School of Life Science and Technology, Institute of Science Tokyo, Tokyo, Japan; Japan Collection of Microorganisms/Microbe Division, RIKEN BioResource Research Center, Ibaraki, Japan

**Keywords:** *Deferribacterota*, protists, termites, *Cryptocercus*, symbiosis, genomics, gut bacteria

## Abstract

Members of the phylum *Deferribacterota* inhabit diverse environments, but their symbiosis with protists has never been reported. We discovered an ectosymbiotic clade of *Deferribacterota* specifically associated with spirotrichonymphid protists in the guts of the termites *Reticulitermes speratus* and *Hodotermopsis sjostedti* and trichonymphid protists in the gut of the wood-feeding cockroach *Cryptocercus punctulatus*. The ectosymbiotic *Deferribacterota* were spiral shaped and attached to 16–91% of the host protist cells. These formed a monophyletic cluster within an uncultured insect gut-associated family-level clade, which is sister to the vertebrate gut-associated family *Mucispirillaceae*. The complete genome of an ectosymbiotic *Deferribacterota* was obtained from a *Trichonympha acuta* cell in a *C. punctulatus* gut and analyzed together with a single-cell amplified genome of another ectosymbiotic *Deferribacterota* associated with *Holomastigotes* sp. in the gut of *R. speratus*. Genome analyses suggest that these *Deferribacterota* ferment monosaccharides and conduct fumarate and oxidative respiration with H_2_ as an electron donor. They thus possibly contribute to the removal of hydrogen and oxygen to protect the fermentative activity of the protist hosts. The ectosymbionts possess reduced signal transduction gene repertoires, implying that the association has provided a relatively stable environment for these bacteria. The ectosymbionts likely possess flagella with an unusually expanded number of flagellin variants up to 40, which may reflect an adaptation to their ectosymbiotic lifestyle. We propose a novel genus, *Termitispirillum*, for these ectosymbionts and a novel family, *Termitispirillaceae*, for the insect-gut clade, under SeqCode. Our findings provide new insights into the ecology and evolution of *Deferribacterota*.

## Introduction

*Deferribacterota* is a phylum of Gram-negative curved rods or spiral-shaped bacteria [1, 2], currently comprising one class (*Deferribacteres*), one order (*Deferribacterales*), and seven families (*Calditerrivibrionaceae*, *Deferribacteraceae*, *Deferrivibrionaceae*, *Flexistipitaceae*, *Geovibrionaceae*, *Mucispirillaceae*, and “*Candidatus* Microvillispirillaceae”) [3–5]. The first five families have been isolated from a wide variety of environments, including deep-sea hydrothermal vents [6], activated sludges [7], terrestrial hot springs [8], and subsurface oil reservoirs [9], whereas the latter two families have exclusively been discovered in animal intestines.

*Mucispirillaceae* inhabit the guts of vertebrates, including rodents [10], chickens [11], and turkeys [12]. *Mucispirillum schaedleri*, the sole cultured species in the family *Mucispirillaceae*, inhabits the guts of rodents and is reportedly involved in host inflammation [10, 13]. “*Candidatus* Microvillispirillaceae” is an uncultured family represented by a single species, “*Candidatus* Rimicarispirillum atlantis”, which occupies the space among microvilli of the ectoperitrophic space in the midgut of the deep-sea hydrothermal vent shrimp genus *Rimicaris* [5, 14, 15]. “*Candidatus* Rimicarispirillum atlantis” can degrade chitin and presumably contributes to the turnover of chitin released during shrimp molt [5, 16]. Members of *Deferribacterota* have also been detected by 16S rRNA amplicon sequencing and metagenomic analyses as minor residents in the guts of insects, including the Auckland tree wētā [17], cockroaches [18, 19], and termites [20–22].

Termites thrive solely on dead plant matter with the aid of their hindgut microbiota [23]. In “lower” termites, i.e., all families except Termitidae (“higher” termites), and their closest relative, the cockroach genus *Cryptocercus*, the digestion of lignocellulose is largely attributable to the gut protists belonging to the phylum *Parabasalia* or *Preaxostyla* [24, 25]. Most of these protists harbor symbiotic prokaryotes in their cytoplasm, nucleoplasm, and/or on the cell surface [26]. It has been suggested that these prokaryotic symbionts contribute to the provision of nitrogenous compounds [27–34] and/or hydrogen removal to promote fermentative activities in the gut [29, 35–38].

During our previous single-cell genomic survey of the bacterial microbiota in the gut of the termite *Reticulitermes speratus* [39], a single-cell amplified genome (SAG), designated MS499-41, was obtained and phylogenetically assigned to *Deferribacterota*. Our preliminary fluorescence in situ hybridization (FISH) analysis implied that the bacterium is an ectosymbiont of a gut protist; however, symbiosis between *Deferribacterota* and protists has never been reported thus far. Thus, in the present study, we aimed to clarify the localization and phylogenetic diversity of *Deferribacterota* in the guts of termites and cockroaches as well as their metabolic capacities and symbiotic roles based on their genome sequences. This study provides evolutionary and functional perspectives on the novel, uncultured *Deferribacterota* clade associated with cellulolytic gut protists.

## Materials and Methods

### Termite and cockroach collection

The Japanese subterranean termite *R. speratus* (family Heterotermitidae) was collected in Tsukuba, Ibaraki Prefecture, and Ogose, Saitama Prefecture in Japan. The damp-wood termite *Hodotermopsis sjostedti* (family Archotermopsidae) was collected in Yakushima, Kagoshima Prefecture in Japan. The wood-feeding cockroach *Cryptocercus punctulatus* was collected in the Black Rock Mountain State Park, Georgia, U.S.A. under permit numbers 192019, 202022, and 222017. The termites and cockroaches were kept with fragments of their nest logs in the laboratory.

### 16S rRNA gene amplicon sequencing and phylogenetic analysis

DNA was extracted from the entire guts of two *C. punctulatus* adults individually using the Power Soil DNA Isolation Kit (MoBio Laboratories). The V3–V4 regions of 16S rRNA (ca. 450 bp) genes were amplified by PCR using the prokaryote-universal primer set Pro341F and Pro805R (Table S1). The library preparation, sequencing on a MiSeq System (Illumina), and amplicon data processing were conducted as previously described [40]. The amplicon sequence variants (ASVs) obtained in this study and in our previous study using 60 termite and eight cockroach species [40] were phylogenetically classified using SINA v1.7.2 [41] with the database SILVA SSURef NR99 release 138.2. The ASVs assigned to *Deferribacterota* were aligned with reference sequences using MAFFT v7 [42]. Ambiguously aligned regions were trimmed using trimAl v1.2 [43]. Maximum likelihood trees were constructed with 1,000 ultrafast bootstrap resampling using IQ-TREE v1.6.12 [44]. The best-fit nucleotide substitution models were selected using ModelFinder [45].

### FISH

The entire guts were removed from worker termites and adult cockroaches, and the gut contents were suspended in Trager’s solution U [46]. The samples were fixed in 4% paraformaldehyde for over 8 hrs at 4°C, rinsed twice with double-distilled water, and placed on MAS-GP glass slides (Matsunami, Japan). FISH targeting 16S rRNA was performed as previously described [47, 48] using oligonucleotide probes labeled with Texas Red or 6-carboxyfluorescein (6FAM). Specific oligonucleotide probes (Table S2) were designed using ARB [49]. The specimens were observed under an Olympus BX51 epifluorescence microscope.

### Collection of protist cells, Sanger sequencing of small subunit (SSU) rRNA genes, and phylogenetic analysis

Termites and cockroaches were dissected, and the gut contents were suspended in sterile Trager’s solution U containing 0.01% bovine serum albumin (BSA). Single protist cells were collected in PCR tubes using a TransferMan NK2 micromanipulator (Eppendorf) equipped with a hand-pulled glass capillary. The collected protist cells were subjected to whole genome amplification (WGA) using a GenomiPhi V2 DNA Amplification Kit (GE Healthcare Life Science). Near full-length 16S and 18S rRNA genes were amplified from the WGA products by PCR using Phusion High-Fidelity DNA polymerase (New England Biolabs). PCR was performed with initial denaturation at 98°C for 3 min, followed by 25 cycles of denaturation at 98L for 30 sec, annealing at 50–70°C for 30 sec, and extension at 72L for 1 min with a final extension at 72L for 10 min. For *Holomastigotes* samples, nested PCR was performed for the amplification of the 18S rRNA genes using primer sets listed in Table S1 and the above program. The PCR products were cloned using a Zero Blunt TOPO PCR Cloning Kit (Thermo Fisher Scientific) after purification using a MonoFas DNA Purification Kit I (ANIMOS, Japan). Sanger sequencing of randomly chosen clones was conducted using a BigDye Terminator v3.1 Cycle Sequencing Kit (Applied Biosystems) on an ABI 3730 Genetic Analyzer. The SSU rRNA gene trees of *Deferribacterota* and their host protists were constructed as described above.

### Genome sequencing and assembly

A SAG of *Deferribacterota* (MS499-41: DRR253682) was obtained in our previous study from the gut content of *R. speratus* using the MDA-in-AGM method and a MiSeq System with a MiSeq Reagent Kit V3 (600 cycles) [39]. In the present study, the MiSeq raw reads were subjected to quality trimming using Cutadapt 3.7 [50] and Prinseq-lite 0.20.4 [51] and were assembled using SPAdes v3.14.0 [52] with the --sc option. Contaminating contigs were identified and removed using the Contig Annotation Tool v5.2.3 [53] and BLASTn searches of the NCBI non-redundant nucleotide database [54]. Only contigs >500 bp were retained.

To acquire the genome sequence of *Deferribacterota* associated with *Trichonympha acuta*, the hindgut content of *C*. *punctulatus* was suspended in sterile Trager’s solution U containing 0.01% BSA, and the membrane portion of a single *T. acuta* cell was separated using a Micro Feather Blade K-715 attached to the micromanipulator and collected in a PCR tube. The sample was subjected to WGA using EquiPhi29 DNA Polymerase (Thermo Fisher Scientific) as previously described [55]. The WGA product was purified using a Zymo Genomic DNA Clean & Concentrator kit (ZYMO Research). Sequencing libraries were prepared using a QIAseq FX DNA Library Kit (QIAGEN) for the MiSeq System and a Native Barcoding Kit 24 V14 for a MinION System (Oxford Nanopore Technologies). Prior to the latter library preparation, debranching, single-strand DNA digestion, purification, and size selection that retained fragments of >2 kbp were conducted as previously described [55]. Sequencing was performed on the MiSeq System as above and on the MinION System with a FLO-MIN114 (R10.4.1) flow cell. After base calling using dorado and quality-filtering using nanoq [56], the MinION reads were assembled using Raven [57]. The resulting circular contig was polished one time using Pilon v1.24 [58] with the quality-filtered MiSeq reads.

The genome quality was assessed using CheckM2 [59]. Circular genomes were visualized using Proksee [60]. The average nucleotide identity (ANI) and average amino acid identity (AAI) were calculated using FastANI [61] and EzAAI [62], respectively. The Skew Index (SkewI), which indicates the degree of GC skew, was calculated using SkewIT [63].

### Phylogenomic analysis

The phylogenomic analysis was conducted using GToTree [64] as the following process. Seventy-four single-copy marker genes conserved among *Bacteria* [64] were extracted from *Deferribacterota* genomes using Prodigal [65] and HMMER [66]. Only genomes that contained ≥50% of the marker genes were retained as references. The deduced amino acid sequences were aligned using MUSCLE [67], trimmed using trimAl v1.2, and concatenated to construct a supermatrix. A maximum likelihood tree was constructed using IQ-TREE as above. The phylogenetic classification was confirmed using the Genome Taxonomy Database (GTDB) [68].

### Functional annotation

The reconstructed genomes were annotated using DFAST [69], KAAS [70], and DIAMOND [71]. Operon structures were predicted using Operon-mapper [72]. CRISPR-Cas systems were identified using CRISPRCasFinder (https://crisprcas.i2bc.paris-saclay.fr/CrisprCasFinder/Index). Protein-coding sequences (CDSs) were assigned to clusters of orthologous groups (COGs) using RPS-BLAST v2.13.0+ [73]. Principal component analysis (PCA) was performed using the sklearn.decomposition module of the Scikit-learn library [74]. Orthologous genes among the *Deferribacterota* genomes were identified using OrthoFinder [75]. Principal coordinate analysis (PCoA) based on orthogroup composition was performed using Phyloseq [76]. The functional annotation of each orthogroup was performed using GhostKOALA [77]. The orthologous genes characteristically associated with a phylogenetic group were identified using Scoary2 [78]. A maximum likelihood tree of the flagellin genes was constructed based on the deduced amino acid sequences with >350 aa as described above.

## Results

### Phylogenetic diversity and relative abundance of *Deferribacterota* in termite and cockroach guts

The phylogenetic diversity and relative abundance of *Deferribacterota* in the guts of termites and cockroaches were examined based on the 16S rRNA gene amplicon data derived from the guts of diverse termite and cockroach species. We identified 72 ASVs assigned to *Deferribacterota* from 47 out of the 68 termite and cockroach species (Fig. S1). Their relative abundance was consistently low (0.00–1.29%). These ASVs formed a monophyletic cluster with sequences derived from other insect guts (Fig. S1). The corresponding 16S rRNA region of the SAG MS499-41 was identical to the sequence of ASV13241 detected in the guts of *R. speratus* (relative abundance: 0.05%) and *Hodotermopsis* sp. (0.06%) (Fig. S1). A near full-length 16S rRNA sequence, RsTz2-092 (TACW01000013), previously obtained from the same *R. speratus* gut sample, showed 99.9% similarity to that of MS499-41. The bacterial species represented by the SAG MS499-41, ASV13241, and RsTz2-092 was designated here as “*Candidatus* Termitispirillum reticulitermitis” (abbreviated as *Te. reticulitermiti*s) and described below.

### Localization, morphology, and infection rate

We designed probe RsTz2-092-190 (Table S2), which is specific to the RsTz2-092 subclade within the insect-gut clade (Fig. S1). FISH using this probe detected long, curved rods or spiral-shaped bacteria, which were specifically attached to the cell surface of the parabasalid protist *Holomastigotes* sp. (family *Spirotrichonymphidae*) in the gut of *R. speratus* (Fig. 1A, B). Morphologically similar bacterial cells were also detected with this probe specifically on the cell surface of *Holomastigotes* sp. (Fig. 1C, D) and *Brugerollina cincta* (*Spirotrichonymphidae*) (Fig. 1E, F) in the gut of *Hodotermopsis sjostedti*. These ectosymbionts appeared to be randomly distributed across the host cell surface (Fig. 1B, D, F) together with spirochaetes, and their cell dimensions were ca. 3–10 µm in length and 0.4 µm in width. Among the cells of the protist host species, 15.8–91.4% harbored ectosymbionts detected with this probe (Table 1).

**Figure 1.**
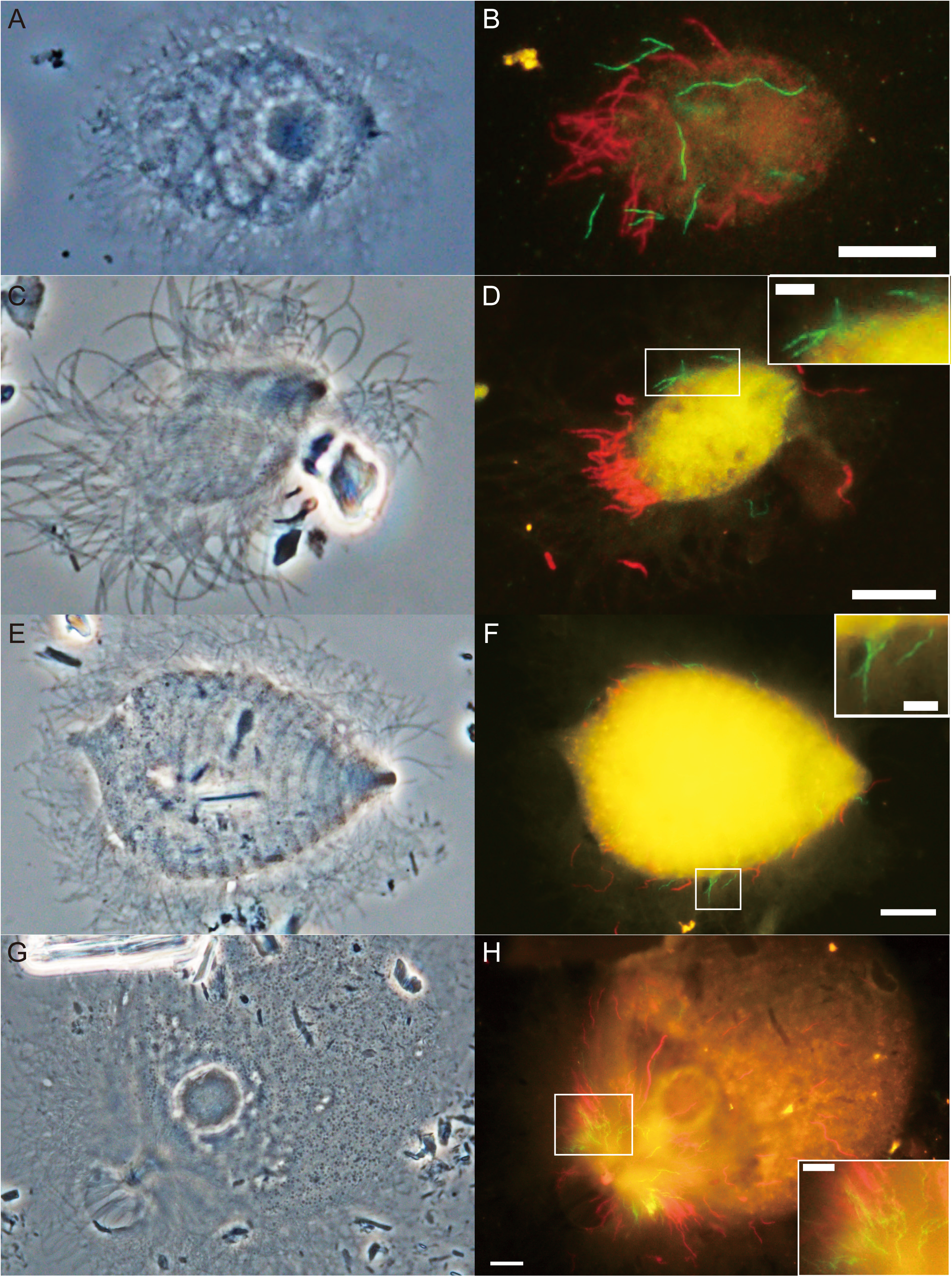
Detection of *Deferribacterota* associated with protists in the guts of termites and a *Cryptocercus* cockroach. Phase-contrast images of (A) *Holomastigotes* sp. from *Reticulitermes speratus*, (C) *Holomastigotes* sp. from *Hodotermopsis sjostedti*, (E) *Brugerollina cincta* from *H*. *sjostedti*, and (G) *Trichonympha lata* in *Cryptocercus punctulatus* are shown. *Deferribacterota* were detected by FISH using probe RsTz2-092-190 labeled with 6FAM (green) (B, D, F) and probe Deferri-term-661 labeled with 6FAM (green) (H), respectively. Spirochaetes were detected using probe Spiro-36 labeled with Texas red (red), and the images were overlayed (B, D, F, H). Bars indicate 10 µm and 5 µm (inset for magnification).

**Table 1.**
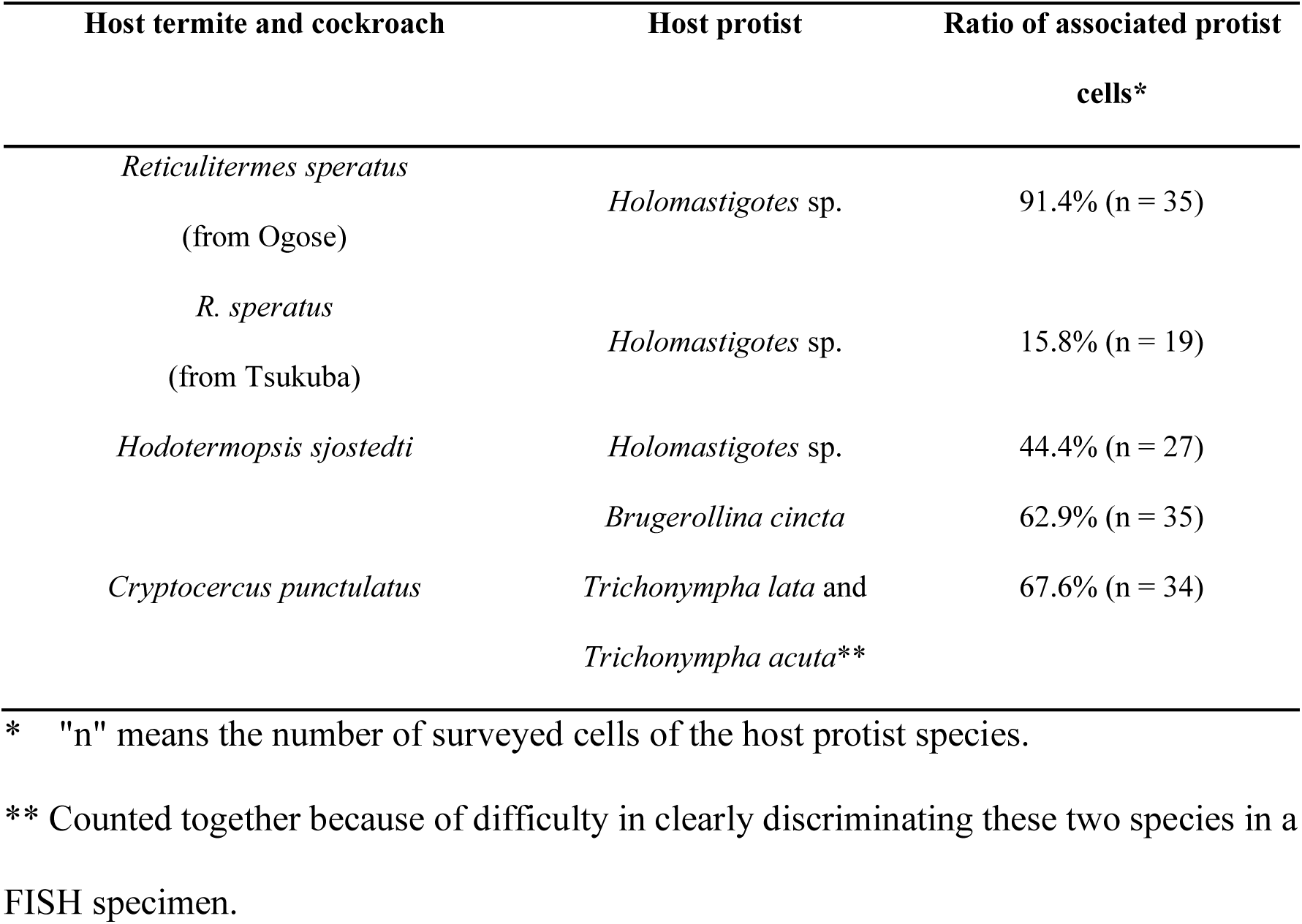
Host protist species and frequency of association with *Deferribacterota*.

We then designed probe Deferri-term-661 (Table S2), which is specific to most members of the insect-gut clade, except for the RsTz2-092 subclade (Fig. S1). When FISH with this probe was performed for the gut content of *C. punctulatus*, spiral cells with dimensions 3–13 µm in length and 0.4 µm in width were detected specifically on the anterior surface area of the parabasalids *Trichonympha lata* (Fig. 1G, H) and *Trichonympha acuta* (Fig. S2A, B) (family *Trichonymphidae*). Of the *T. lata* and *T. acuta* cells in a *C. punctulatus* gut, 67.6% harbored the ectosymbiotic *Deferribacterota* (Table 1). Ectosymbiotic spirochaetes were also detected on the same host protist cells by FISH using a spirochaete-specific probe (Figs. 1H and S2B; Table S2). No specific signals were detected in the gut contents of *R. speratus* and *H. sjostedti* with probe Deferri-term-661.

### Phylogenetic analysis based on near full-length 16S rRNA genes

We collected single cells of these host protist species, i.e., *Holomastigotes* spp., *B. cincta*, *T. acuta*, and *T. lata*, based on their morphological characteristics and performed WGA. Near full-length 18S rRNA and 16S rRNA genes were amplified by PCR and the products were cloned and subjected to Sanger sequencing. We confirmed the taxonomic assignment of the host protist species based on the 18S rRNA genes (Fig. S3A, B). The 16S rRNA genes of *Deferribacterota* formed a monophyletic cluster of sequences within the insect-gut clade (Fig. 2A). The clone RsHol6-1, obtained from *Holomastigotes* sp. in an *R. speratus* gut, showed 99.9% sequence similarity to the clone RsTz2-092. The clones HsHol1-2 from *Holomastigotes* sp. and HsBr1-3 from *B. cincta* in the gut of *Hodotermopsis sjostedti* were identical and showed 98.7% similarity to RsHol6-1 and RsTz2-092 (Fig. 2A). The clones CpT15-2 and CpT06-9 were obtained from *T. acuta* and *T. lata*, respectively, in the gut of *C. punctulatus*. These two sequences shared 97.9% identity with each other (Fig. 2A) and showed ≥99.3% similarities to ASV149, which was detected in one *C. punctulatus* gut (relative abundance: 0.21%) (Fig. S1).

**Figure 2.**
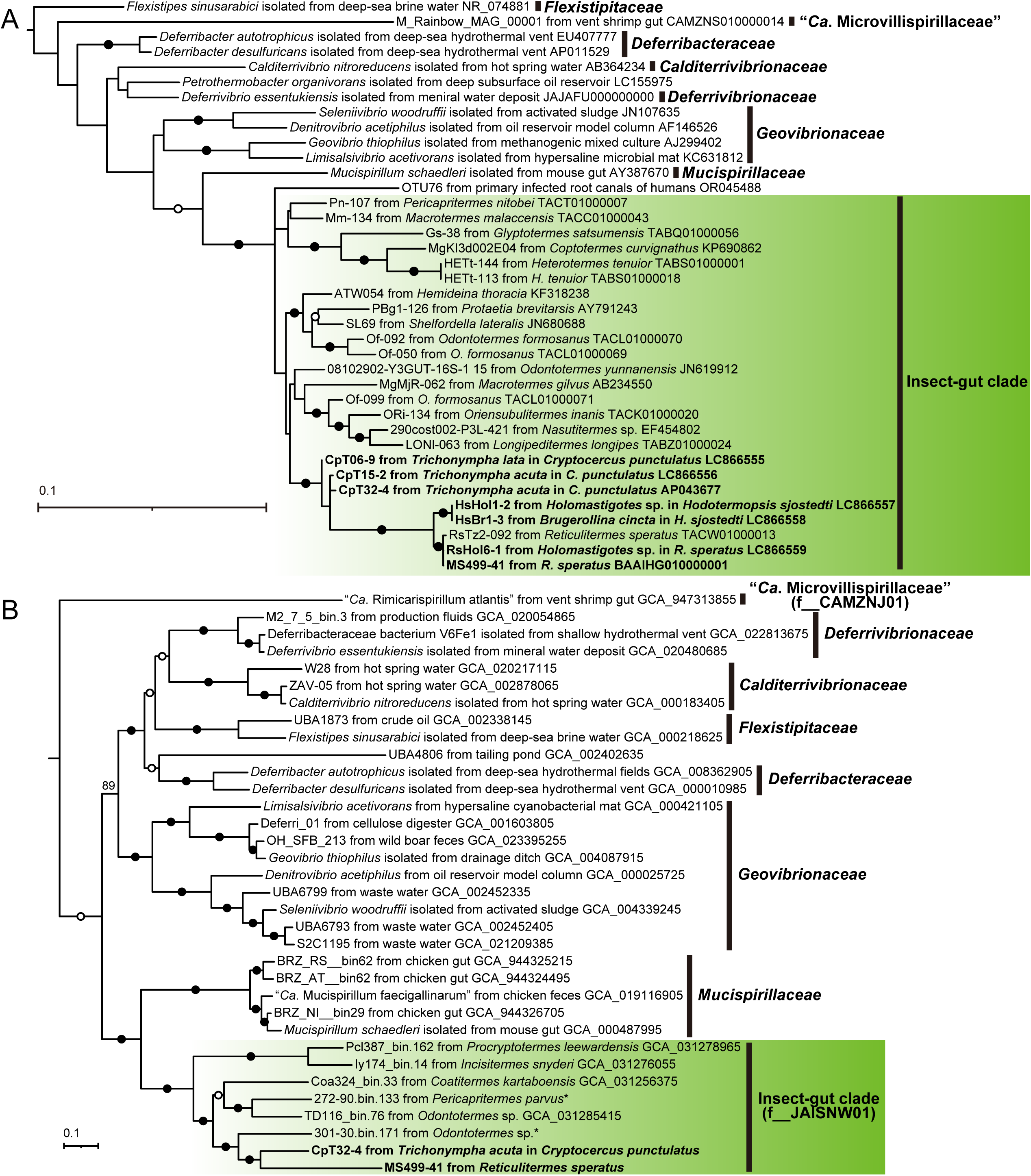
Phylogenetic positions of the ectosymbiotic *Deferribacterota*. (A) 16S rRNA gene tree constructed using 993 nucleotide sites with the GTR+F+I+G4 substitution model and an outgroup (AB729138). (B) Phylogenomic tree constructed using 12,478 amino acid sites of concatenated, partitioned 74 marker genes, with substitution models selected for each part and an outgroup (GCF_000469585 and GCF_000177635). Family-level taxonomic codes according to GTDB are indicated in parentheses. Filled and open circles indicate nodes with ultrafast bootstrap value ≥95% and ≥80% in panel A, and 100% and ≥90% in panel B, respectively. Asterisks indicate sequences obtained by Arora et al. (2022).

### General features of *Termitispirillum* genomes and phylogenomics

In addition to the SAG MS499-41 (1,578,178 bp, 79% estimated completeness, 44% GC content) of *Te. reticulitermitis* from the gut of *R. speratus*, a circular chromosome (CpT32-4: 2,151,577 bp, 40% GC content) was reconstructed from a single *T. acuta* cell sample (Table 2). The 16S rRNA gene sequence of CpT32-4 shared 98.0% and 98.4% identity with CpT15-2 and CpT06-9, respectively (Fig. 2A). The bacterium represented by the genome CpT32-4 was designated here as *Termitispirillum cryptocerci* and described below under SeqCode. The total sequence length and GC content of the *Te. reticulitermitis* and *Te. cryptocerci* genomes were similar to those of other insect-gut clade species (Fig. S4). The GC skew pattern of the *Te. cryptocerci* genome is distinct from other genomes of *Deferribacterota*; the SkewI of *Te. cryptocerci* is only 0.176, whereas the others range 0.733–0.914 (Fig. S5). The *Te. cryptocerci* genome encodes CRISPR and associated genes (Table S3), whereas they were not found in the *Te. reticulitermitis* genome.

**Table 2.** General genome features of MS499-41, CpT32-4, and related *Deferribacterota*.

|  | MS499-41 | CpT32-4 | *Mucispirillum*<br>*schaedleri* | "*Rimicarispirillum*<br>*atlantis*" |
| --- | --- | --- | --- | --- |
| Family | Insect-gut clade | Insect-gut clade | *Mucispirillaceae* | "Microvillispirillaceae" |
| Localization | Ectosymbiont of<br>*Holomastigotes* sp. | Ectosymbiont of<br>*T. acuta* | Mucus layer in the<br>mouse gut | Space among microvilli<br>in the shrimp gut |
| Total length (Mb) | 1.58 | 2.15 | 2.35 | 1.34 |
| Contigs | 254 | 1 (circular) | 1 (circular) | 49 |
| Completeness (%)* | 78.6 | Complete | Complete | 90.7 |
| Contamination (%)* | 0.5 | - | - | 0.0 |
| GC content (%) | 44.1 | 39.7 | 31.1 | 47.4 |
| CDS | 1,363 | 1,931 | 2,145 | 1,072 |
| Coding density (%) | 78.8 | 89.3 | 87.9 | 84.5 |
| rRNA | 4 | 6 | 9 | 3 |
| tRNA | 36 | 40 | 42 | 33 |

\* Estimated using CheckM2.

The genomes of *Te. reticulitermitis* and *Te. cryptocerci* were most closely related to metagenome-assembled genomes (MAGs) from the guts of higher termites [79, 80], which generally do not harbor cellulolytic gut protists. These and two additional MAGs from lower termites (Pcl387_bin.162 and Ly174_bin.14) [33] formed a monophyletic cluster (Fig. 2B). This clade corresponds to the uncultured family “JAISNW01” in the GTDB and is phylogenetically sister to the vertebrate-gut inhabitants *Mucispirillaceae* (Fig. 2B). The ANI and AAI values between the *Te. reticulitermitis* and *Te. cryptocerci* genomes were <70% and 62.0%, respectively (Fig. S6A, B). Thus, these two bacteria are distinct species but in the same genus according to the proposed criterion of genus-level similarity (ANI ≤ 95%; AAI ≥ 60–85%) [61, 81].

### Metabolic capacities of Te. cryptocerci and Te. reticulitermitis

#### Energy conservation and carbon metabolism

The *Te. cryptocerci* genome contains a complete set of genes for glycolysis and gluconeogenesis (Fig. 3A, B). *Termitispirillum cryptocerci* likely imports monosaccharides and sugar acids, such as fructose, xylose, mannose, N-acetylglucosamine, and uronate, via phosphotransferase systems (PTS), ABC transporters, or transporters of major facilitator superfamilies (MFS). It also likely imports and utilizes glycerol, lactate, citrate, and unidentified carboxylates. Citrate can be metabolized to acetate and oxaloacetate by the action of citrate lyase (CitDEF) (Fig. 3A). It is predicted that pyruvate is fermented to acetate, ethanol, lactate, and H_2_ using monomeric iron hydrogenase B1/B3 and electron-bifurcating iron hydrogenase HydABC (Fig. 3A, B). The *Te. reticulitermitis* genome lacks two genes in the glycolysis/gluconeogenesis pathway and all genes encoding the transporters for sugars and other carbon sources, except for those transporting uronates and carboxylates (Figs 3A, B, and S7), possibly due to its low completeness (Table 2). Thus, the main carbon source of *Te. reticulitermitis* remains unclear. No genes encoding hydrolytic enzymes for extracellular polysaccharides were identified in either genome.

**Figure 3.**
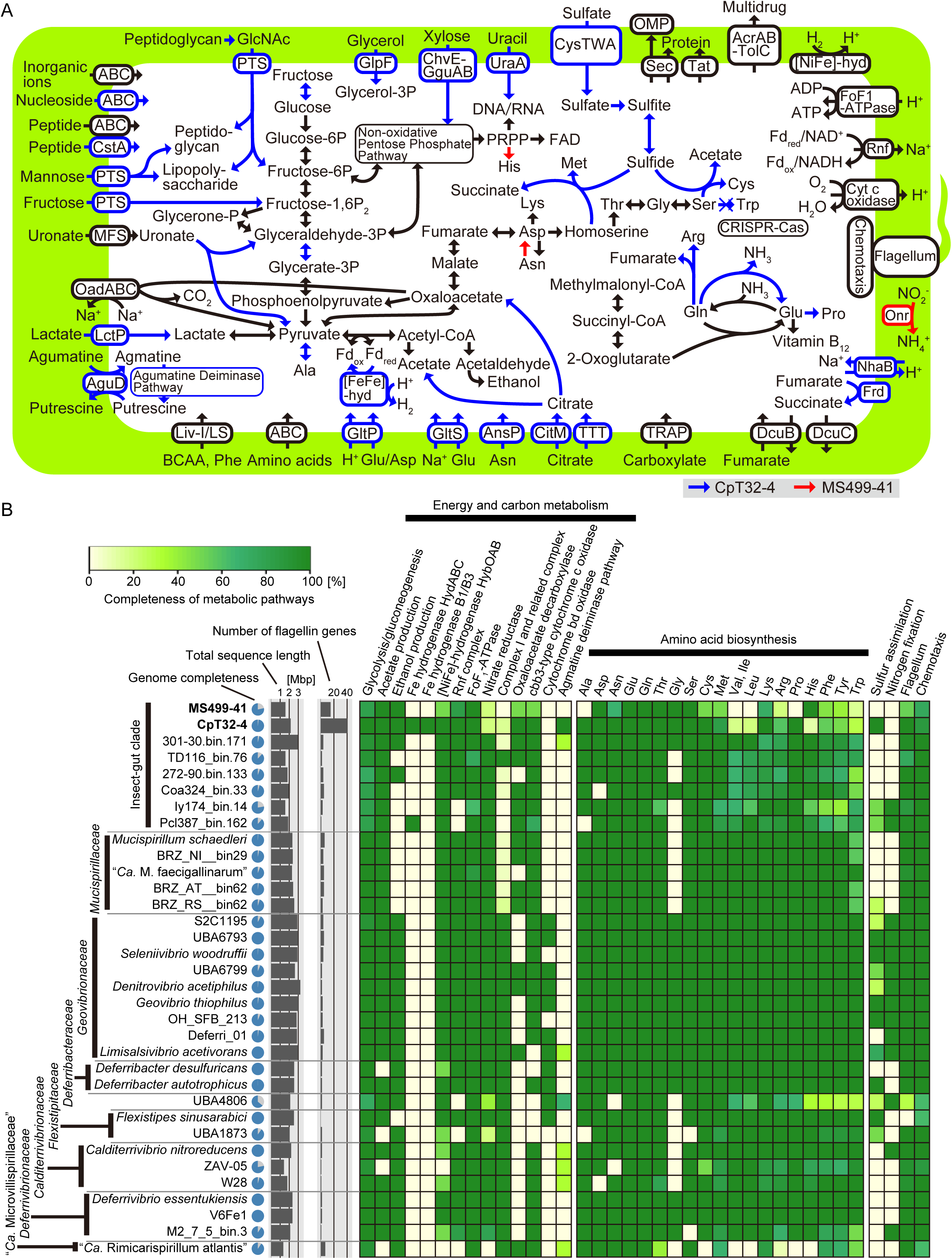
Predicted metabolic capacities of *Termitispirillum cryptocerci* (CpT32-4) and “*Candidatus* Termitispirillum reticulitermitis” (MS499-41). (A) Outline of predicted metabolic pathways based on genome sequences. Pathways detected exclusively in *Te. cryptocerci* or *Te. reticulitermitis* are indicated in blue and red, respectively. (B) Comparisons of repertoires of genes and pathways among members of *Deferribacterota*. The completeness of pathways is indicated in a heatmap. Estimated genome completeness, total sequence length, and the number of flagellin genes are also shown.

In addition to substrate-level phosphorylation through glycolysis and oxidation to acetate, *Te. cryptocerci* has the potential to generate ATP via the agmatine deiminase pathway with the agmatine/putrescine antiporter AguD [82] (Fig. 3A). The *Te. cryptocerci* genome is equipped with several systems to generate H^+^/Na^+^ membrane potential to produce ATP via FoF_1_-type ATPase. The genome encodes the membrane-bound [NiFe]-hydrogenase HybOABC, the Rnf complex, Na^+^-transporting oxaloacetate/methylmalonyl-CoA decarboxylase (OadABC), and the H^+^/Na^+^ antiporter NhaB. It also encodes fumarate reductase (FrdABCD) with the fumarate/succinate antiporter DcuB and the succinate exporter DcuC, and in addition, *cbb*_3_-type cytochrome *c* oxidase (Fig. 3A). All these components except for NhaB and FrdABCD were identified in the *Te. reticulitermitis* genome. Thus, the ectosymbionts likely use H_2_ as an electron donor and fumarate and oxygen as electron acceptors for respiration. Considering the very low concentration of oxygen in the hindguts of termites and cockroaches, except for their peripheral regions [83], the *cbb*_3_-type cytochrome *c* oxidase may function not only in ATP synthesis via microaerobic respiration but also in oxygen removal to protect anaerobic reactions. The *Te. reticulitermitis* genome encodes a gene for periplasmic octaheme nitrite reductase (Onr) (Figs. 3A and S7), which possibly contributes to the detoxification of nitrite by dissimilatory reduction to ammonium, as suggested in *Desulfovibrio* species [84]. The genomes of these ectosymbionts contain several other genes for oxygen tolerance (Fig. S7).

#### Nitrogen metabolism

No genes encoding a nitrogenase were identified in the *Te. cryptocerci* and *Te. reticulitermitis* genomes. Both genomes encode several transporters for amino acids and di/oligo-peptides, which are likely the main nitrogen sources (Fig. 3A). The *Te. cryptocerci* genome also encodes transporters for nucleosides and uracil although both genomes apparently can de novo synthesize nucleotides. Both genomes lack the biosynthetic pathways for branched-chain amino acids, and only the *aroE*, *aroK, aroA, and aroC* genes were identified in the pathway for aromatic amino acids. Although an operon for the tryptophan synthase genes *trpA* and *trpB* was found in the *Te. cryptocerci* genome, both are pseudogenized. *Termitispirillum cryptocerci* cannot synthesize histidine, whereas *Te. reticulitermitis* possesses its complete biosynthetic pathway. These biosynthetic pathways, absent in *Te. cryptocerci* and/or *Te. reticulitermitis*, are typically retained by most other *Deferribacterota* (Fig. 3B). In the *Te. cryptocerci* genome, the biosynthetic pathways for riboflavin, tetrahydrofolate, cobalamin, and heme are complete, whereas the biosynthesis of pantothenate/coenzyme A (CoA), nicotinamide adenine dinucleotide (NAD), biotin, pyridoxin 5’-phosphate, and menaquinone requires precursors (Fig. S7).

#### Sulfur metabolism

*Termitispirillum cryptocerci* potentially takes up sulfate, which is subsequently reduced to sulfite via adenosine 5’-phosphosulfate (APS) and 3’-phosphoadenosine 5’-phosphosulfate (PAPS). It is predicted that sulfite is reduced to sulfide by the action of assimilatory sulfite reductase (Sir) in cooperation with the heterodisulfide reductase complex HdrABC-FlxABCD (Fig. S8A, B) [85]. Sulfides can be assimilated into methionine and cysteine. No genes involved in sulfur assimilation were detected in the *Te. reticulitermitis* genome (Fig. 3A, B).

#### Cell wall, secretion systems, and motility

Both genomes encode the biosynthetic pathways for peptidoglycan and lipopolysaccharides and also contain the gene sets encoding proteins for flagellar assembly and chemotaxis, similar to the cases in other *Deferribacterota* [3, 5, 6, 13] (Fig. 3A, B). Genes for the secretory (Sec) system and the twin arginine translocation (Tat) pathway, which facilitate protein transport, were present, whereas no other secretion systems were identified (Fig. 3A).

#### Comparison of genomic compositions with other members of Deferribacterota

PCA based on the relative abundance of COG categories exhibited the clustering of the gut-inhabiting clades, i.e., the insect-gut clade, *Mucispirillaceae*, and “*Ca*. Microvillispirillaceae” (Fig. 4). Among the COG categories, a decrease in the ratio and number of genes in category [T] (signal transduction mechanisms) was prominent in *Te. cryptocerci* and *Te. reticulitermitis* (Figs 4 and S9A, B). In PCoA based on the composition of all orthologs (Fig. S10A, B), the genomes of the insect-gut clade formed a cluster and were separated from those of other animal-gut inhabitants.

**Figure 4.**
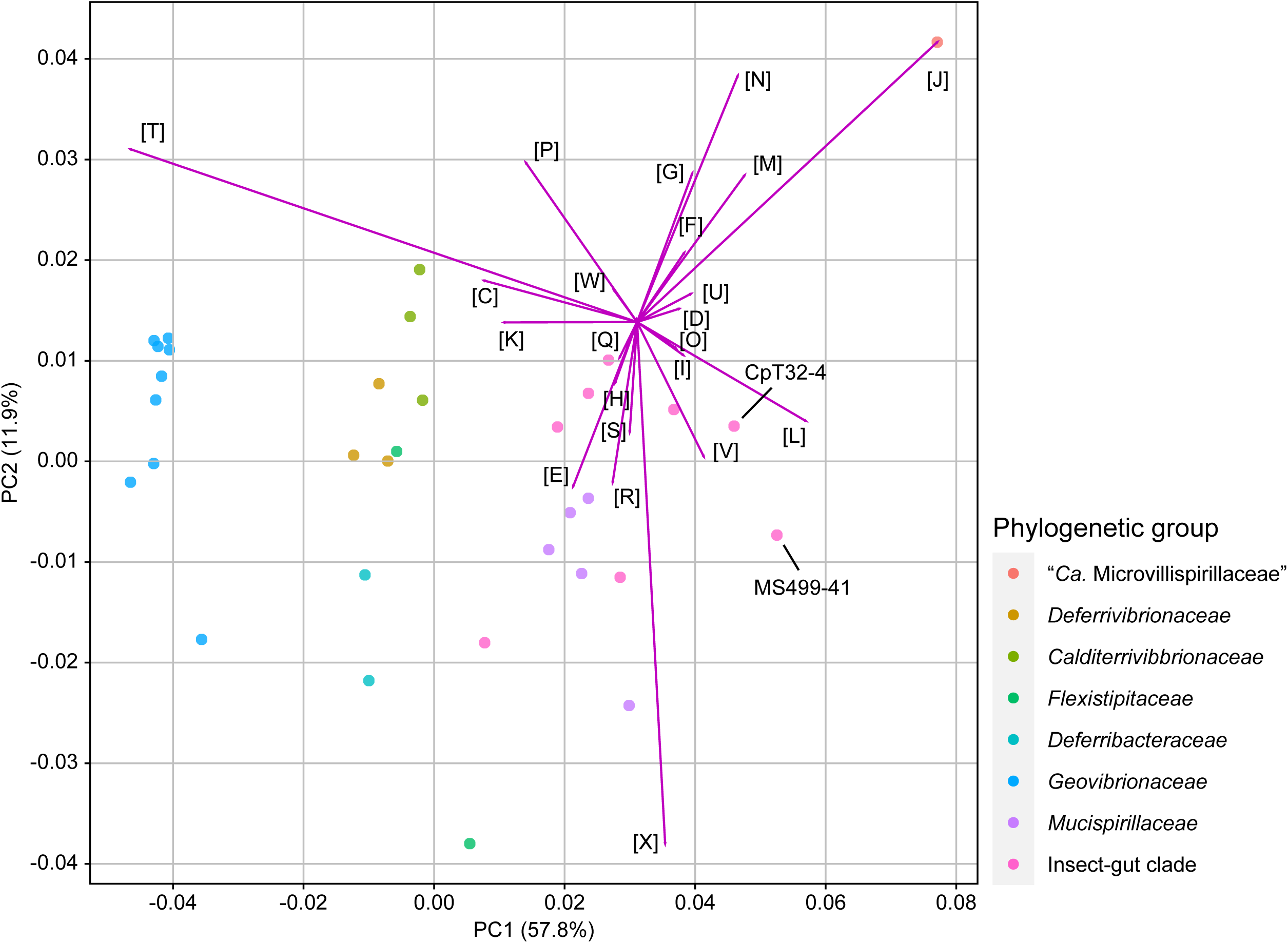
Comparation of genomic contents among members of *Deferribacterota*, including *Termitispirillum cryptocerci* (CpT32-4) and “*Candidatus* Termitispirillum reticulitermitis” (MS499-41). Principal component analysis based on the relative abundance of the clusters of orthologous groups (COGs) was performed. The COG categories: [C] Energy production and conversion, [D] Cell cycle control, cell division, chromosome partitioning, [E] Amino acid transport and metabolism, [F] Nucleotide transport and metabolism, [G] Carbohydrate transport and metabolism, [H] Coenzyme transport and metabolism, [I] Lipid transport and metabolism, [J] Translation, ribosomal structure and biogenesis, [K] Transcription, [L] Replication, recombination and repair, [M] Cell wall/membrane/envelope biogenesis, [N] Cell motility, [O] Posttranslational modification, protein turnover, chaperones, [P] Inorganic ion transport and metabolism, [Q] Secondary metabolites biosynthesis, transport and catabolism, [R] General function prediction only, [S] Function unknown, [T] Signal transduction mechanisms, [U] Intracellular trafficking, secretion, and vesicular transport, [V] Defense mechanisms, [W] Extracellular structures, [X] Mobilome: prophages, transposons.

Most genomes of the insect-gut clade and *Mucispirillaceae* lack components for nitrogen fixation and NADH dehydrogenase complex I (Fig. 3B). Among the *Deferribacterota*, *Te. cryptocerci* exceptionally has capacity to use diverse carbon and energy sources (Fig. S7) and to generate H_2_ as a fermentation product (Fig. 3B). In addition, the agmatine deiminase pathway is almost unique to *Te. cryptocerci* in *Deferribacterota*, and its ability to assimilate sulfate is rare among the gut inhabitants. The biosynthetic capacity for amino acids and cofactors is reduced in the insect-gut clade and especially in the ectosymbionts *Te. cryptocerci* and *Te. reticulitermitis*. This tendency is also conspicuous in the shrimp-gut symbiont “*Ca.* Rimicarispirillum atlantis” in “*Ca*. Microvillispirillaceae” (Figs 3B and S7). *Termitispirillum cryptocerci* and *Te. reticulitermitis* possess an unusually high copy number of flagellin genes compared with other *Deferribacterota* species (*Te. cryptocerci*: 40 copies, *Te. reticulitermitis*: 15 copies, others: 0–7 copies) (Figs 3B and S11).

## Discussion

In this study, we discovered the first examples of ectosymbiosis between *Deferribacterota* and protists. The ectosymbiotic *Deferribacterota* are associated with specific parabasalid protist species in the guts of termites and *Cryptocercus* cockroaches, although not all cells of the host protist species are colonized. The monophyly of these bacteria suggests that their ability to establish ectosymbiosis with protist cells may have evolved in the gut microbiota of the common ancestor of termites and *Cryptocercus*, i.e., before ∼150 million years [86, 87], after the insect-gut clade diverged from *Mucispirillaceae*, a vertebrate gut-associated family that includes *Mucispirillum schaedleri*, a member of the Schaedler flora [88].

The *Deferribacterota* detected in the guts of higher termites lacking cellulolytic gut protists are most likely free-swimming bacteria with flagella (Fig. 3B). The ectosymbionts *Te. cryptocerci* and *Te. reticulitermitis* possess genes encoding proteins for flagellar assembly and chemotaxis; therefore, these bacteria also swim, possibly targeting host protists, and attach to the host cell surfaces. The expansion of the number of flagellin gene variants in the genomes of *Te. cryptocerci* and *Te. reticulitermitis*, up to 40, is noteworthy. In many bacteria, flagella act not only for swimming but also for other roles such as adhesion and colonization [89]. It has been suggested that such multiple functions of flagella are associated with the structural variations of flagellin [90]. In addition, in pathogenic bacteria, shifts in the expression of flagellin variants to avoid or confuse recognition of their flagella by the host cell were previously reported [89]. We thus hypothesize that the large repertoire of flagellin variants may be used for multiple functions, such as swimming and adhesion, and possibly also to avoid recognition and elimination by the protist host.

Most protist cells in the guts of termites and *Cryptocercus* cockroaches harbor ectosymbiotic bacteria of diverse lineages, which likely play various important roles in the multilayered symbiotic system [26]. Our genome analyses indicated that *Te. cryptocerci* and *Te. reticulitermitis* are not able to fix dinitrogen. In addition, they have limited capacity to synthesize amino acids (Fig. 3B), whereas their ability to synthesize several cofactors, such as riboflavin, cobalamin, and tetrahydrofolate, may contribute to the nutrition of the protist and termite hosts (Fig. S7). Although *Te. cryptocerci* has the potential to assimilate diverse carbon sources, including plant-derived monosaccharides and uronate, it does not possess secretory enzymes for the hydrolysis of polysaccharides (Fig. 3A).

As *Te. cryptocerci* and *Te. reticulitermitis* possess the ability to consume H_2_ as an energy source and use O_2_ as an electron acceptor, these ectosymbionts possibly play a role in the removal of these molecules to promote the fermentative activities of their anaerobic protist hosts. This is in concordance with the localization of *Te. cryptocerci* confined to the anterior part of *Trichonympha* cells, where hydrogenosomes are densely localized [91], although this restriction may alternatively reflect higher phagocytotic activity in the posterior region. This localization pattern was previously observed in H_2_-oxidizing ectosymbiotic *Desulfovibrio* associated with *Trichonympha* in termite guts [36, 37]. Thus, the ectosymbiotic *Deferribacterota* are potentially beneficial to their protist hosts, yet they are not consistently maintained across all host cells. This suggests that they are not essential for the host survival.

The reduction in the number of signal transduction-related genes, with no prominent genome size reduction, was previously found in several obligate ectosymbionts of protists in the guts of termites and *Cryptocercus* cockroaches [37]. This trait is shared by *Te. cryptocerci* and *Te. reticulitermitis*; it is implied that these bacteria almost entirely live as ectosymbionts, with free-living or infectious stages before colonizing a host occurring only transiently. The unusual GC skew pattern of *Te. cryptocerci* suggests a non-canonical chromosome replication system and/or massive genome rearrangements, as previously found in endosymbiotic bacteria of termite-gut protists [27, 55, 92] (Fig. S5). However, as no complete genomes except for *Te. cryptocerci* were obtained, it remains unclear whether this GC skew pattern is a common trait in the insect-gut clade or attributable to the ectosymbiosis. Additional genome sequences of ectosymbiotic *Deferribacterota* and ecological characterization of other members of the insect-gut clade will further provide insights into the evolution of ectosymbiosis from free-swimming gut bacteria.

We here describe the novel genus *Termitispirillum* (represented by the type species *Te. cryptocerci*) and a novel family, *Termitispirillaceae*, for the insect-gut clade, under the Code of Nomenclature of Prokaryotes Described from Sequence Data (SeqCode) [93]. We also propose the novel species “*Candidatus* Termitispirillum reticulitermitis” and “*Candidatus* Termitispirillum hodotermopsidis”, which lack high-quality genomic information, as recommended for uncultured bacteria under the International Code of Nomenclature of Prokaryotes [94].

### Description of *Termitispirillaceae* fam. nov

*Termitispirillaceae* (Ter.mi.ti.spi.ril.la’ce.ae. N.L. neut. n. *Termitispirillum*, the type genus of the family; -*aceae*, suffix denoting a family; N.L. fem. pl. n. *Termitispirillaceae*, the family of *Termitispirillum*). Members of this family are uncultured and inhabit the guts of insects, including termites and *Cryptocercus* cockroaches. This family is a sister clade of the family *Mucispirillaceae* in the phylum *Deferribacterota*. The type genus of this family is *Termitispirillum*.

### Description of *Termitispirillum* gen. nov

*Termitispirillum* (Ter.mi.ti.spi.ril’lum. L. masc. n. *termes*, termite; L. neut. dim. n. *spirillum*, a small spiral; N.L. neut. n. *Termitispirillum*, a spiral bacterium from termites). Members of this genus are uncultured and spiral shaped or curved long rods that specifically attach to parabasalid protists in the guts of termites and *Cryptocercus* cockroaches. Cells measure 3–13 µm in length and 0.4 µm in width. Genomic analyses indicated that these bacteria possess a Gram-negative-type cell wall and chemoheterotrophic metabolism with fermentation and respiratory pathways. Motility was inferred from the presence of genes involved in flagellar assembly and chemotaxis. The type species is *Termitispirillum cryptocerci* with its genome sequence as the type material.

### Description of Termitispirillum cryptocerci sp. nov

*Termitispirillum cryptocerci* (cryp.to.cer’ci. N.L. gen. n. *cryptocerci*, of *Cryptocercus*, referring to the host cockroach genus). The bacterium specifically attaches to the cell surfaces of *Trichonympha acuta* and *Trichonympha lata* in the gut of *Cryptocercus punctulatus*. The cell dimensions are 3–13 µm by 0.4 µm. This assignment is based on the complete genome sequence (AP043677) and detection by FISH using the oligonucleotide probe Deferri-term-661 (Table S2). This species corresponds to CpT32-4.

### Description of “*Candidatus* Termitispirillum reticulitermitis” sp. nov

*Termitispirillum reticulitermitis* (re.ti.cu.li.ter’mi.tis. N.L. gen. masc. n. *reticulitermitis*, of *Reticulitermes*, referring to the host termite genus). The bacterium specifically attaches to the cell surface of *Holomastigotes* sp. in the gut of *Reticulitermes speratus*. The cell dimensions are 3–10 µm by 0.4 µm. This assignment is based on the 16S rRNA gene sequence (LC866559) and specific detection by FISH using the oligonucleotide probe RsTz2-092-190 (Table S2). The single-cell genome MS499-41 (BAAIHG010000001–BAAIHG010000254) was obtained. This species includes the 16S rRNA phylotypes RsTz2-092 and RsHol6-1.

### Description of “*Candidatus* Termitispirillum hodotermopsidis” sp. nov

*Termitispirillum hodotermopsidis* (ho.do.ter.mop’si.dis. N.L. fem. n. *hodotermopsidis*, of *Hodotermopsis*, referring to the host termite genus). The bacterium specifically attaches to the cell surfaces of *Holomastigotes* sp. and *Brugerollina cincta* in the gut of *Hodotermopsis sjostedti*. The cell dimensions are 3–10 µm by 0.4 µm. This assignment is based on the 16S rRNA gene sequences (LC866557, LC866558) and specific detection by FISH using the oligonucleotide probe RsTz2-092-190 (Table S2). This species includes the 16S rRNA phylotypes HsHol1-2 and HsBr1-3.

## Acknowledgements

We are grateful to Gaku Tokuda, Aram Mikaelyan, and Melbert Schwartz for cooperation in cockroach collection. We also thank Shu Onouchi, Hirokazu Kuwahara, and Kazuki Takahashi for assisting with experiments and Takumi Murakami for handling sequence data. Sanger sequencing was performed in the RIKEN CBS and the Biomaterial Analysis Center in the Tokyo Institute of Technology.

## Author contributions

Naoya Maruoka: conceptualization, methodology, investigation, formal analysis, data curation, visualization, resource, writing – original draft, funding acquisition. Rinpei Kudo: investigation, data curation. Katsura Igai: investigation, data curation. Masahiro Yuki: methodology, investigation, data curation, resource. Michiru Shimizu: methodology, investigation, data curation, resource. Moriya Ohkuma: resource, writing – review and editing, funding acquisition. Yuichi Hongoh: conceptualization, methodology, resource, formal analysis, writing – original draft, writing – review and editing, funding acquisition, project administration, supervision.

## Data availability

The genome sequences obtained in this study are available under the BioProject PRJDB20420 in DDBJ. The accession numbers of the SSU rRNA genes appears under DDBJ accession numbers LC866555–64 and LC875758.

