## Supplemental Tables for "Discovery and genomics of H_2_-oxidizing/O_2_-reducing *Deferribacterota* ectosymbiotic with protists in the guts of termites and a *Cryptocercus* cockroach"

**Table S1.** PCR primers used in this study.

| Primer name | Target gene | Primer sequence (5′→3′) | Reference |
|---|---|---|---|
| Pro341F | Prokaryote 16S rRNA gene (V3–V4 region) | CCTACGGGNBGCASCAG | [1] |
| Pro805R | Prokaryote 16S rRNA gene (V3–V4 region) | GACTACNVGGGTATCTAATCC | [1] |
| 27Fmix | *Bacteria* 16S rRNA gene (near full-length) | AGRGTTTGATYMTGGCTCAG | [2] |
| 1492Rmix | *Bacteria* 16S rRNA gene (near full-length) | GGHTACCTTGTTACGACTT | [2] |
| SpiroF1 | *Parabasalia* 18S rRNA gene (outer) | ATACTTGGTCGATCCTGCCAAGG | [3] |
| SpiroR1 | *Parabasalia* 18S rRNA gene (outer) | TGATCCAACGGCAGGTTCMCCTAC | [3] |
| GGF | *Parabasalia* 18S rRNA gene (inner) | CTTCGGTCATAGATTAAGCCATGC | [4] |
| GGR | *Parabasalia* 18S rRNA gene (inner) | CCTTGTTACGACTTCTCCTTCCTC | [4] |
| E23F3 | *Eukarya* 18S rRNA gene (near full-length) | ACYTGGTTGATYCTGCC | [5] |
| E1511R4 | *Eukarya* 18S rRNA gene (near full-length) | CWDCBGCAGGTTCWCCWAC | [5] |

**Table S2.** FISH probes used in this study.

| Probe name | Target taxon | Probe sequence (5′→3′) | Reference |
|---|---|---|---|
| RsTz2-092-190 | RsTz2-092 clade | GCCTTTCTTGCTACACCA | This study |
| Deferri-term-661 | Insect gut-clade | TATTCCGCATTCCTCTCC | This study |
| Spiro-36 | *Spirochaetales* | CTTAAGACGCGCCGCCAG | [6] |
| EUB338 | Most bacteria | GCTGCCTCCCGTAGGAGT | [7] |

* Except for RsTz2-092 and its close relatives.

**Table S3.** Repeat and spacer sequences of CRISPR in the CpT32-4 genome.

| | Sequence (5′→3′) | Position |
|---|---|---|
| Repeat | CGCACCTTTCACGGGTGCGTGGATTGAAAT | |
| Spacer 1 | ACCGATTGGCAAAAGCTACAACTGCCCGCAGTGC | 144,837–144,870 |
| Spacer 2 | ACGTAAAATCGAGCCTTTATCCGATACGAGCATGA | 144,901–144935 |
| Spacer 3 | ACCGCTGGCGGACACGCTATCACAGAGCCGTACATT | 144,966–145,001 |
| Spacer 4 | ATAGATTTAGGTGTAGTTATAAAGCCATACACTAG | 145,032–145,066 |
| Spacer 5 | ATTTTCGCACAATGATCATTTACAGCCTCGACAGG | 145,097–145,131 |
| Spacer 6 | TGTAGTAAAGTGTTAGCAGTTTACTAACAGAGGT | 145,162–145,195 |
| Spacer 7 | TATTGAGCATGAGCCACTAATCCCGACCCTTATA | 145,226–145,259 |
| Spacer 8 | AGCTTATTGGATCAAAAAAAGACCGTGTTGAAA | 145,290–145,322 |
| Spacer 9 | ACAAAGGCCAATAGGCGAGCACAAGTGTGCAAGAG | 145,353–145,387 |
| Spacer 10 | GTATAGGCAGTGATCGTTACCTAGCTGCAACTCCAG | 145,418–145,453 |
| Spacer 11 | TGGCTAATCGTAAAATATACGATGCTGACAAACT | 145,484–145,517 |
| Spacer 12 | TCAAATTATGTGCCGATCATGACACATAAAAAGT | 145,548–145,581 |
| Spacer 13 | TTATTTTATACTCCATCTCTGCCAGCCGCCGCTT | 145,612–145,645 |
| Spacer 14 | ATATTGTTACTTTTAGTGCTAACAAACACACCAA | 145,674–145,709 |
| Spacer 15 | TGAATAGGTAGAAAGGGTGTGTATTATGGAAATG | 145,740–145,773 |
| Spacer 16 | TCAGCTCTGCGCCATCAACGCCCAGCTCTTTCAG | 145,804–145,837 |
| Spacer 17 | TATTAATTTATCCTTCCTTATTTTTATGCTTCGG | 145,868–145,901 |

**References to Supplementary Materials**
