## Supplemental Figures for "Discovery and genomics of H_2_-oxidizing/O_2_-reducing *Deferribacterota* ectosymbiotic with protists in the guts of termites and a *Cryptocercus* cockroach"

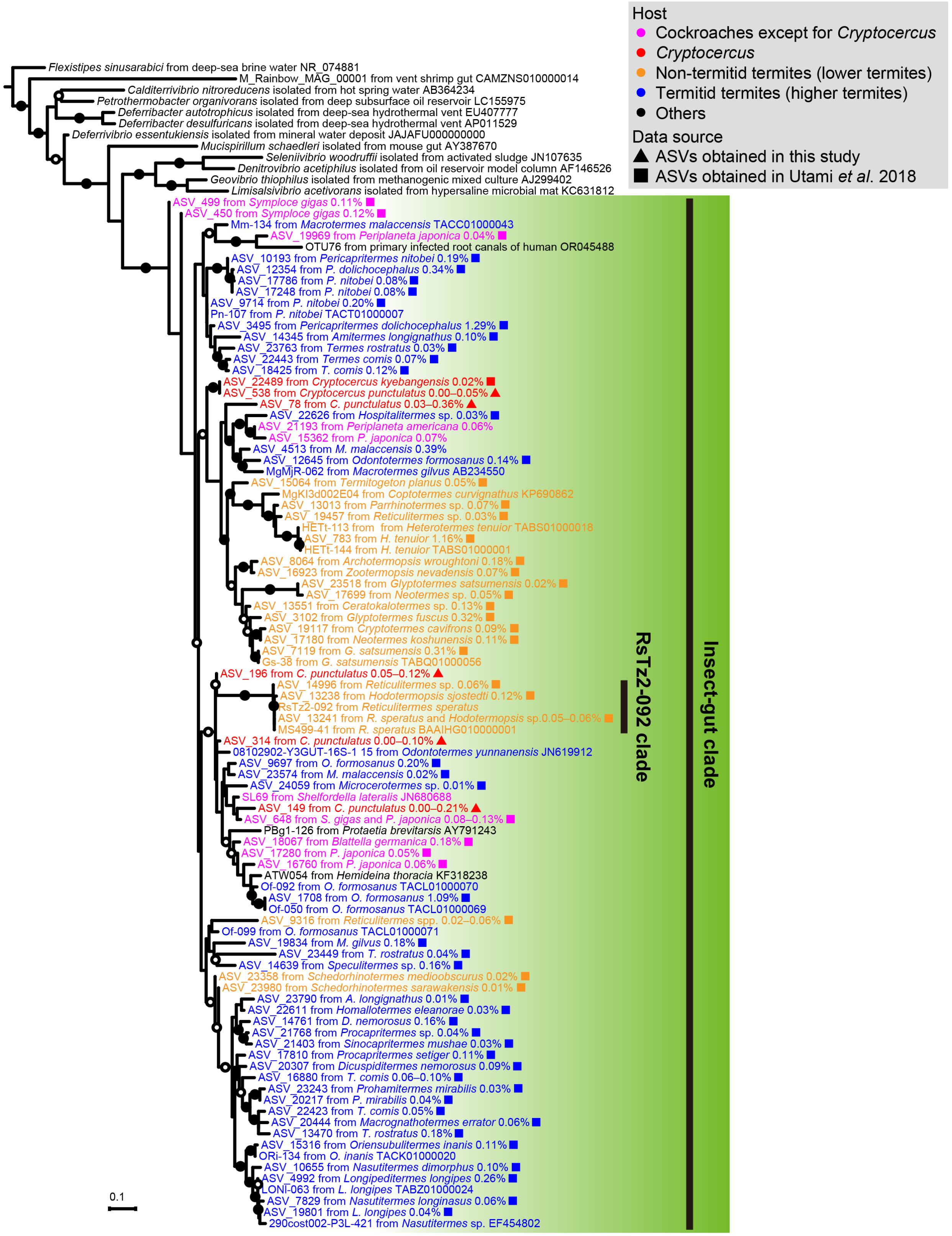


**Figure S1.** Phylogenetic diversity of *Deferribacterota* in termite and cockroach guts based on the 16S rRNA gene V3–V4 region. A maximum-likelihood tree was constructed using 409 nucleotide positions with the TIM3e+I+G4 substitution model and an outgroup (AB729138). Filled and open circles indicate nodes with ultrafast bootstrap value ≥95% and ≥80%, respectively.

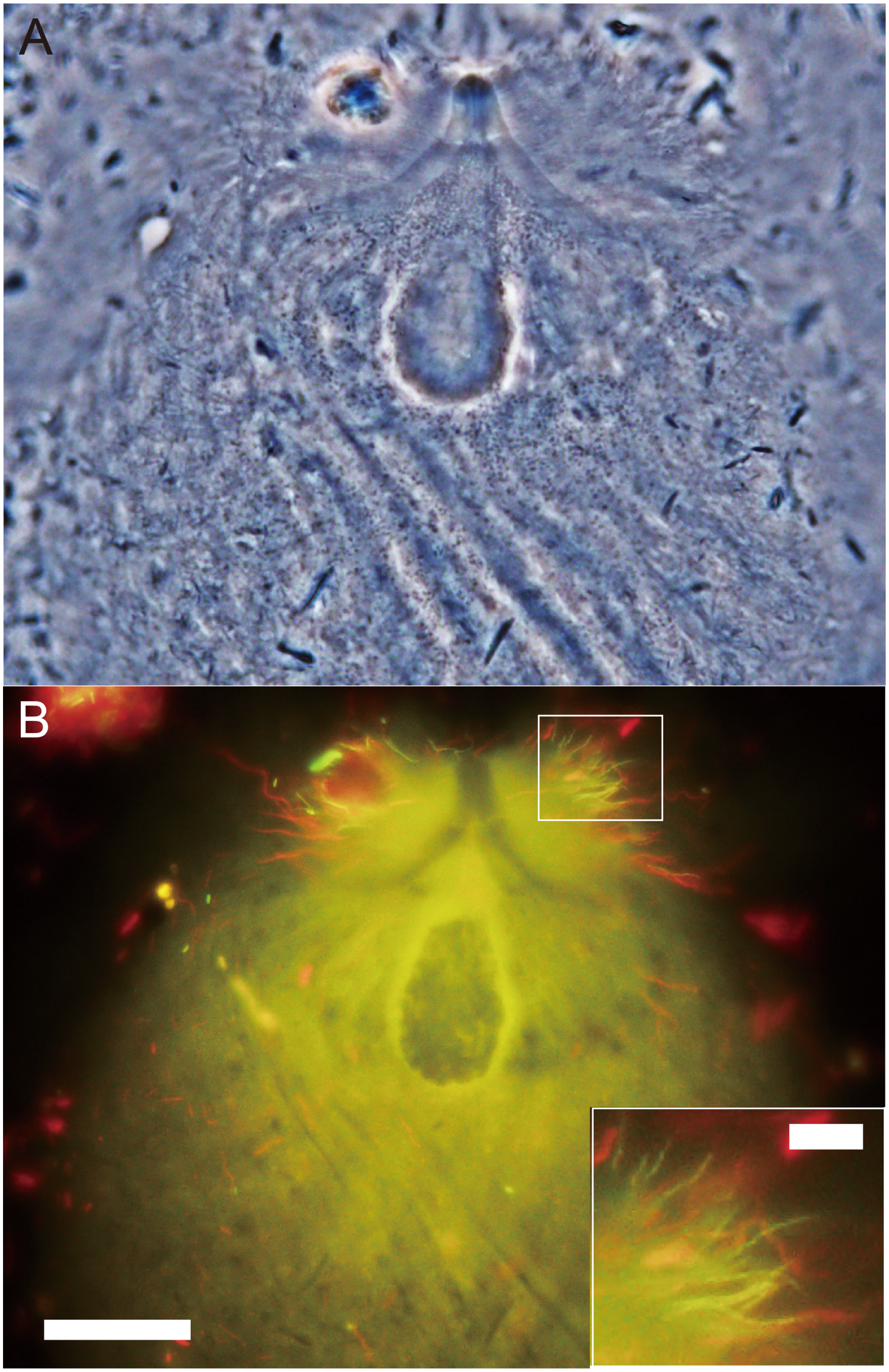


**Figure S2.** Detection of *Deferribacterota* associated with protists in the gut of *Cryptocercus punctulatus*. (A) Phase-contrast image of *Trichonympha acuta*. (B) Epifluorescence image. FISH using probes Deferri-term-661 labeled with 6FAM (green) and EUB338 labeled with Texas red (red) was performed. The FISH images were overlayed; spiral, rod-shaped cells of *Deferribacterota* were detected with a merged, yellowish color. Magnified image is shown in inset. Bar indicates 50 μm and 10 μm (inset).

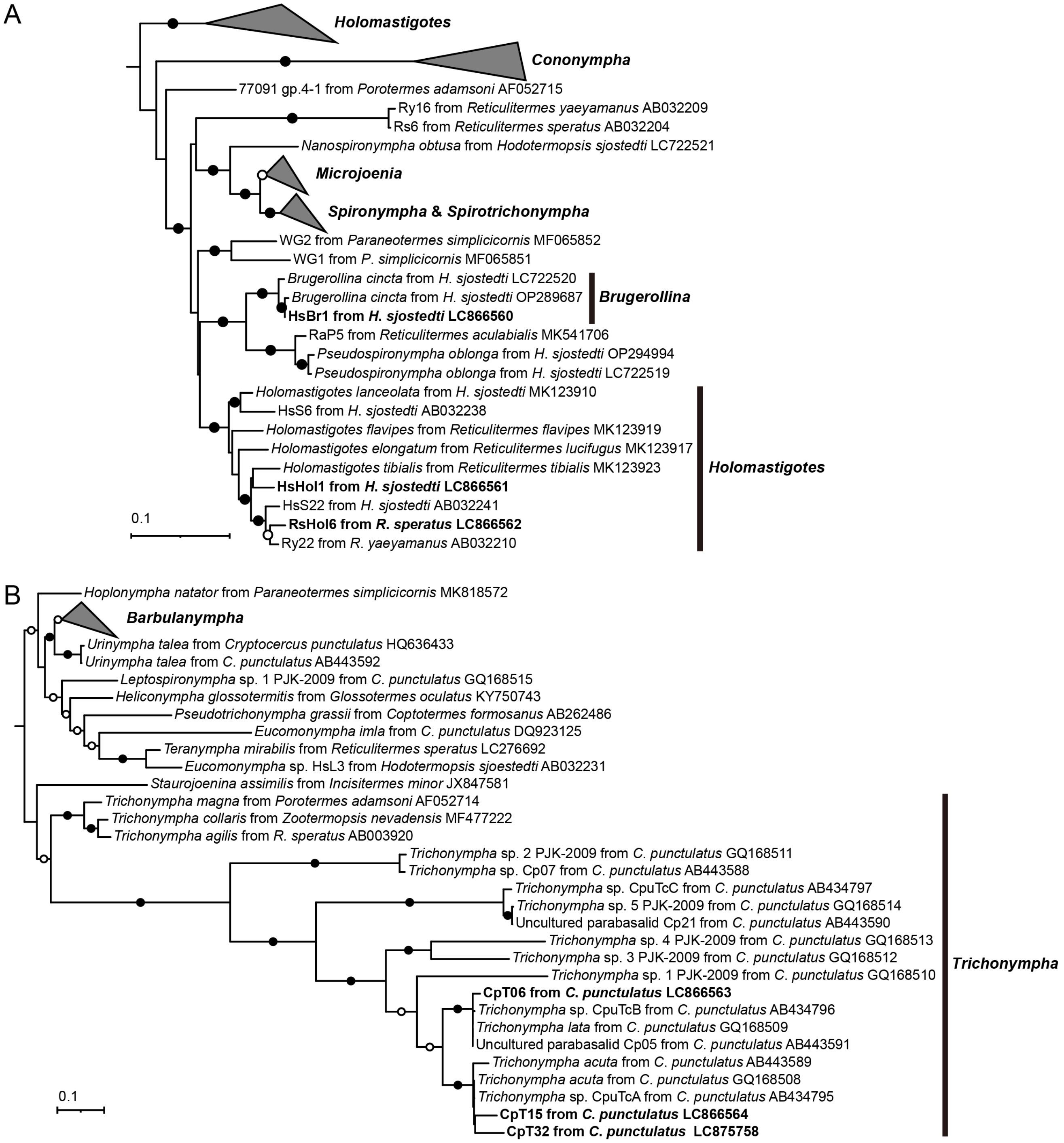


**Figure S3.** Phylogenetic positions of host protists based on the 18S rRNA gene. Maximum likelihood trees of (A) *Spirotrichonymphida* constructed using 1,364 nucleotide sites with the TVMe+I+G4 substitution model and an outgroup (AY055799, AY319278, GQ254638, and GQ254637) and (B) *Trichonymphida* constructed using 1,276 nucleotide sites with the GTR+F+I+G4 substitution model and an outgroup (AY338476 and JN619423) are shown. The sequences obtained in this study are shown in bold. Filled and open circles indicate ultrafast bootstrap value ≥ 95% and ≥80%, respectively.

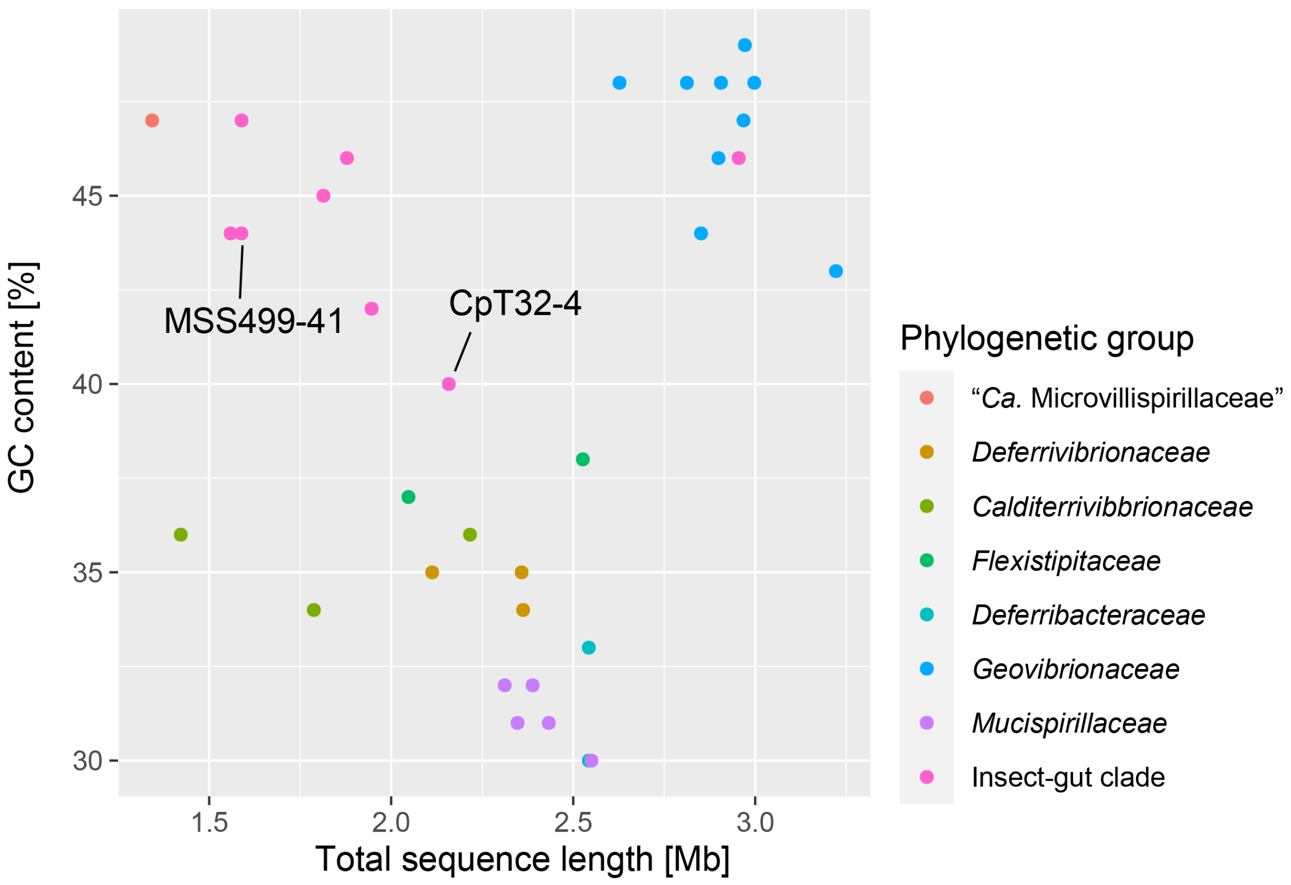


**Figure S4.** Comparisons of total sequence length and GC content of *Deferribacterota* genomes, which were used in the phylogenomic analysis in Fig. 2B. Genomes with an estimated completeness of <75% were excluded. The genomes of *Termitispirillum cryptocerci* (CpT32-4) and "*Candidatus* Termitispirillum reticulitermitis" (MS499-41) are indicated.

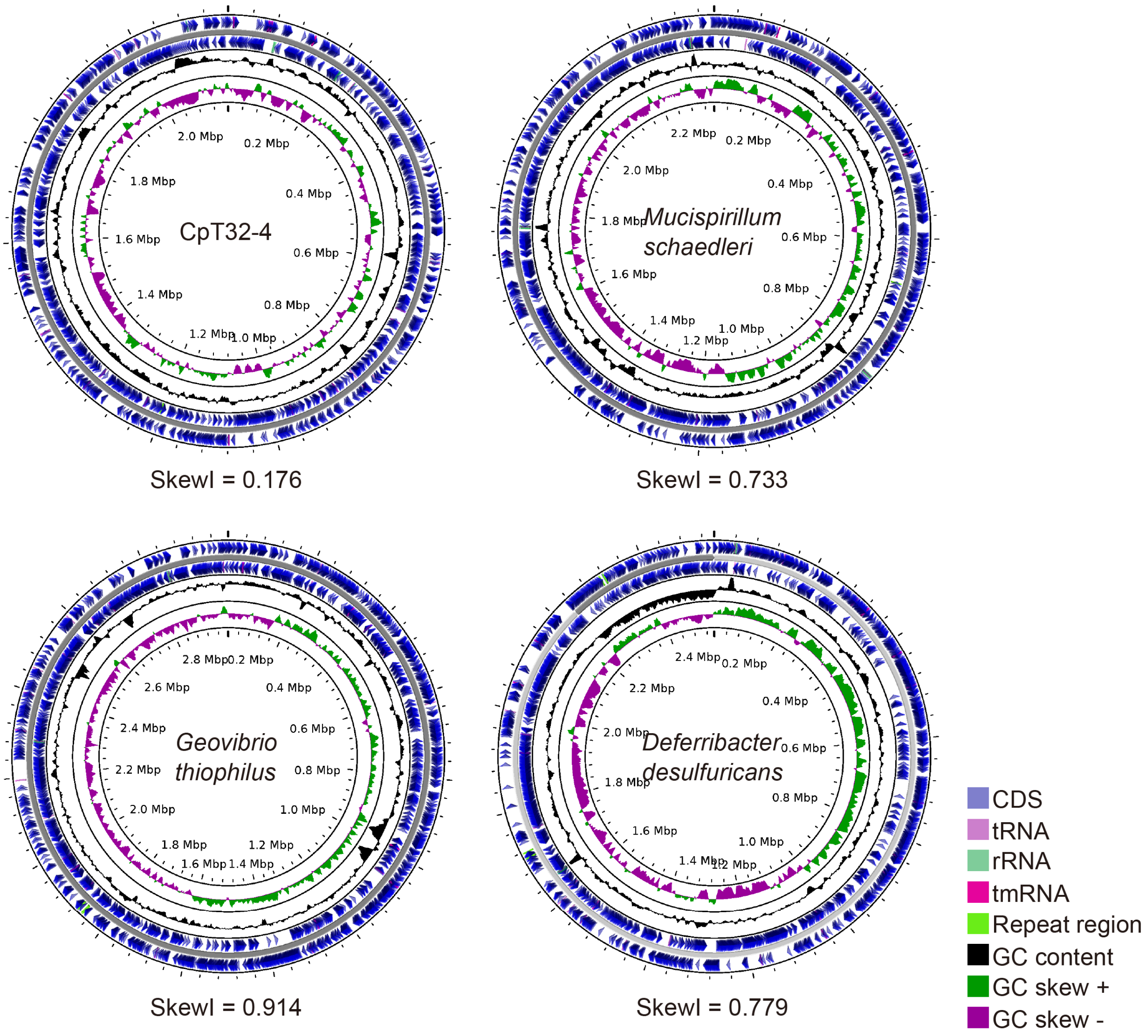


**Figure S5.** Circular maps of complete chromosomal genomes of four *Deferribacterota* species, including *Termitispirillum cryptocerci* (CpT32-4). Position 1 is set at the initial nucleotide site of the *dnaA* gene. SkewI values are shown.

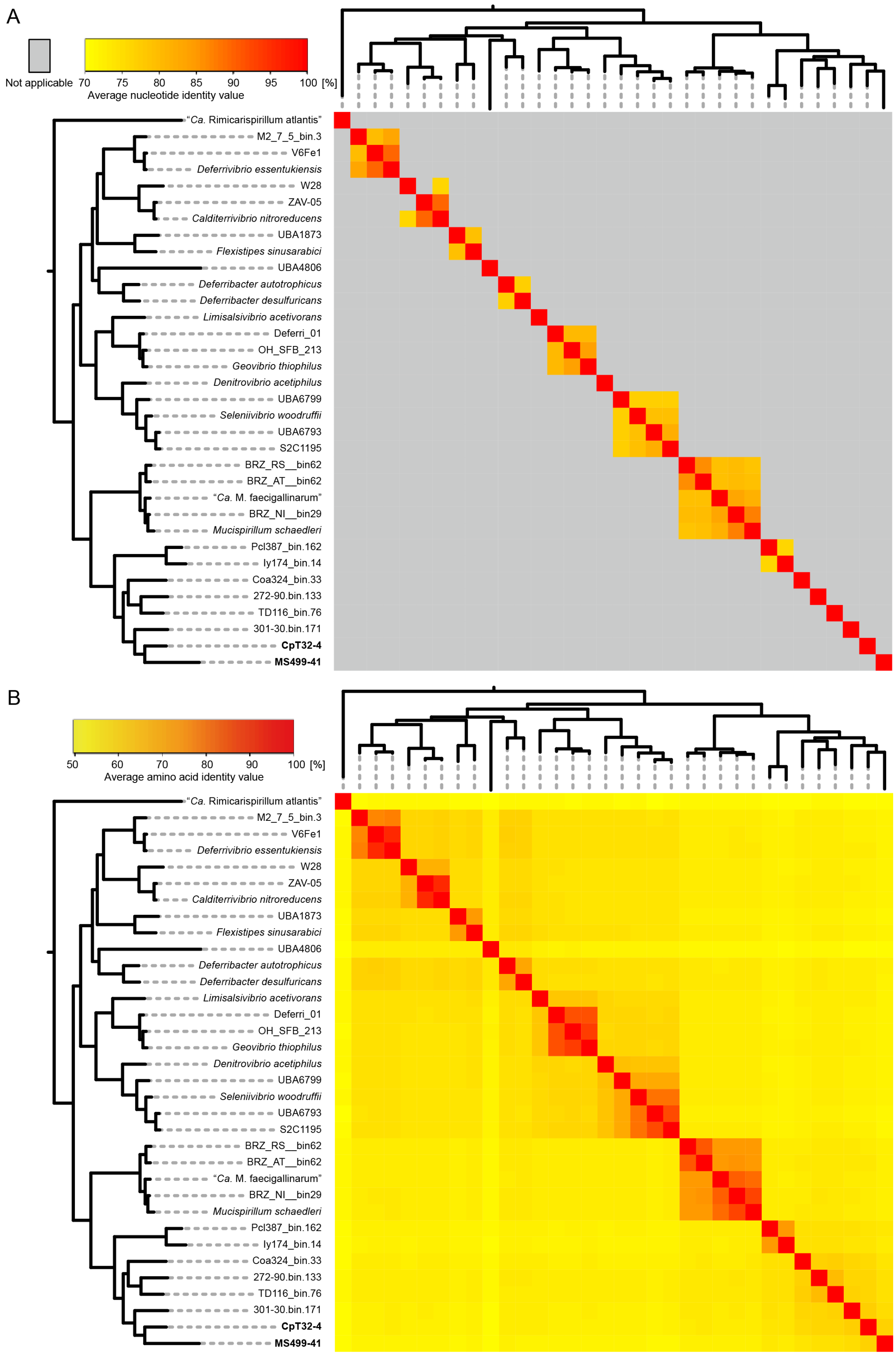


**Figure S6.** Pairwise sequence identities between *Deferribacterota* genomes, including *Termitispirillum cryptocerci* (CpT32-4) and "*Candidatus* Termitispirillum reticulitermitis" (MS499-41). (A) Average nucleotide identities and (B) average amino acid identities are presented as heatmaps.

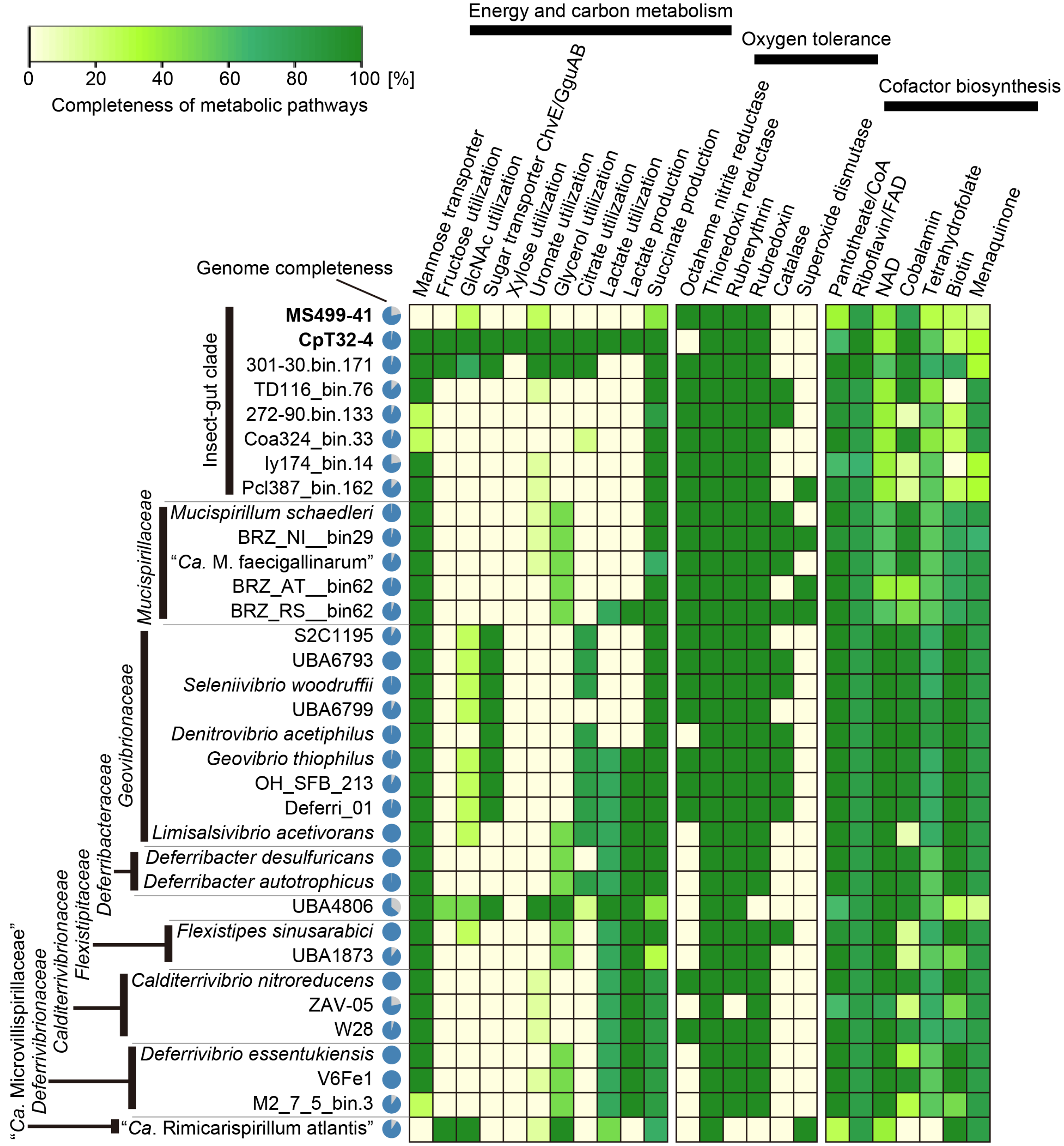


**Figure S7.** Presence or absence of genes or completeness of metabolic pathways in *Deferribacterota*, including *Termitispirillum cryptocerci* (CpT32-4) and "*Candidatus* Termitispirillum reticulitermitis" (MS499-41).

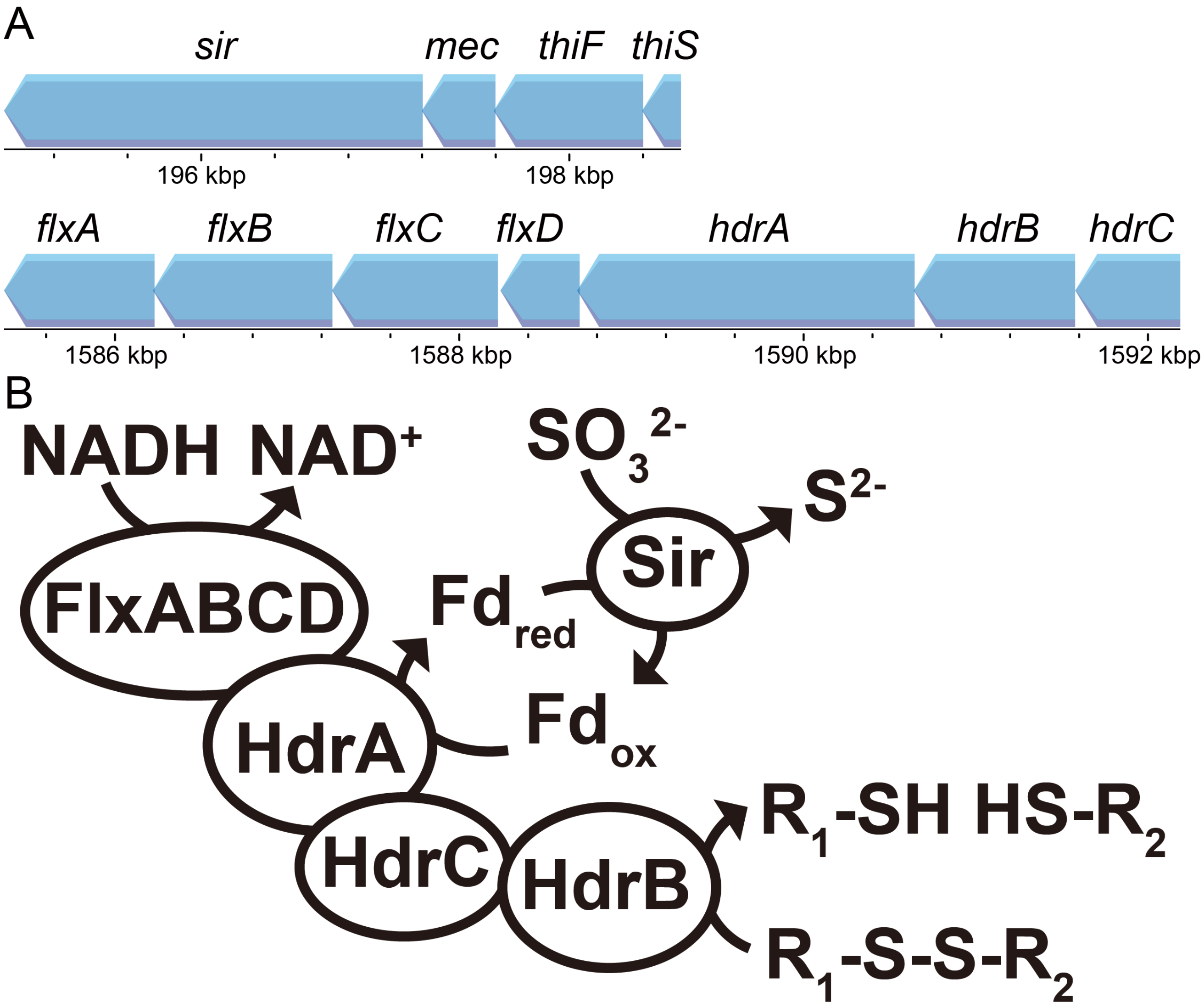


**Figure S8.** Genes putatively involved in sulfite reduction in *Termitispirillum cryptocerci* (CpT32-4). (A) Loci of genes encoding sulfite reductase (*sir*), [CysO sulfur-carrier protein]-S-L-cysteine hydrolase (*mec*), sulfur carrier protein ThiS adenylyltransferase (*thiF*), sulfur carrier protein (*thiS*), NADH dehydrogenase complex (*flxABCD*), and heterodisulfide reductase complex (*hdrABC*). Arrows indicate the direction of transcription. (B) Predicted scheme of sulfite reduction driven by a flavin-based electron bifurcation.

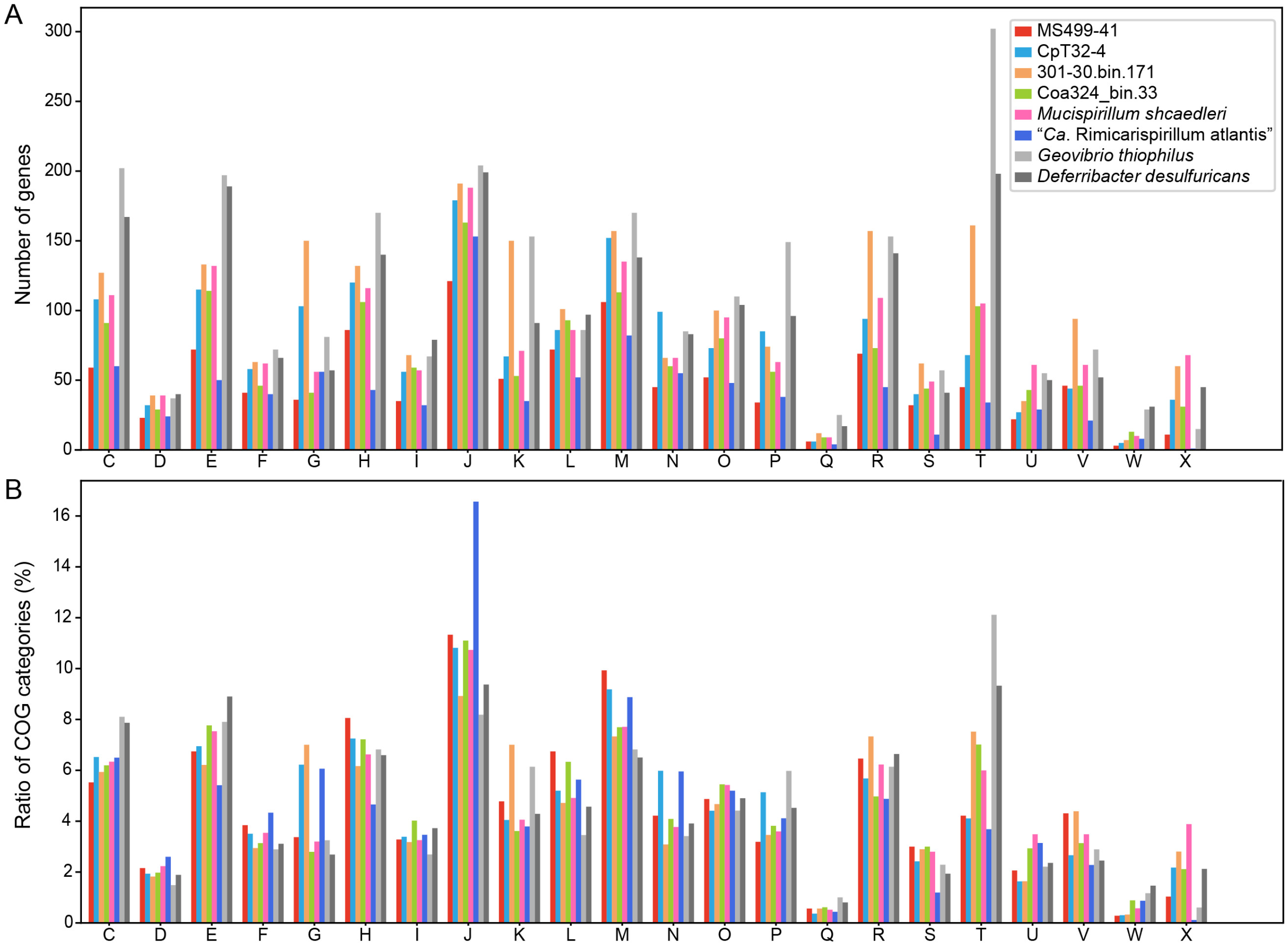


**Figure S9.** Comparation of genomic contents based on the clusters of orthologous genes (COGs) among representative species of *Deferribacterota*, including *Termitispirillum cryptocerci* (CpT32-4) and "*Candidatus* Termitispirillum reticulitermitis" (MS499-41. (A) Total number and (B) ratio of genes per COG category in each *Deferribacterota* species. The COG categories analyzed are: [C] Energy production and conversion, [D] Cell cycle control, cell division, chromosome partitioning, [E] Amino acid transport and metabolism, [F] Nucleotide transport and metabolism, [G] Carbohydrate transport and metabolism, [H] Coenzyme transport and metabolism, [I] Lipid transport and metabolism, [J] Translation, ribosomal structure and biogenesis, [K] Transcription, [L] Replication, recombination and repair, [M] Cell wall/membrane/envelope biogenesis, [N] Cell motility, [O] Posttranslational modification, protein turnover, chaperones, [P] Inorganic ion transport and metabolism, [Q] Secondary metabolites biosynthesis, transport and catabolism, [R] General function prediction only, [S] Function unknown, [T] Signal transduction mechanisms, [U] Intracellular trafficking, secretion, and vesicular transport, [V] Defense mechanisms, [W] Extracellular structures, [X] Mobilome: prophages, transposons.

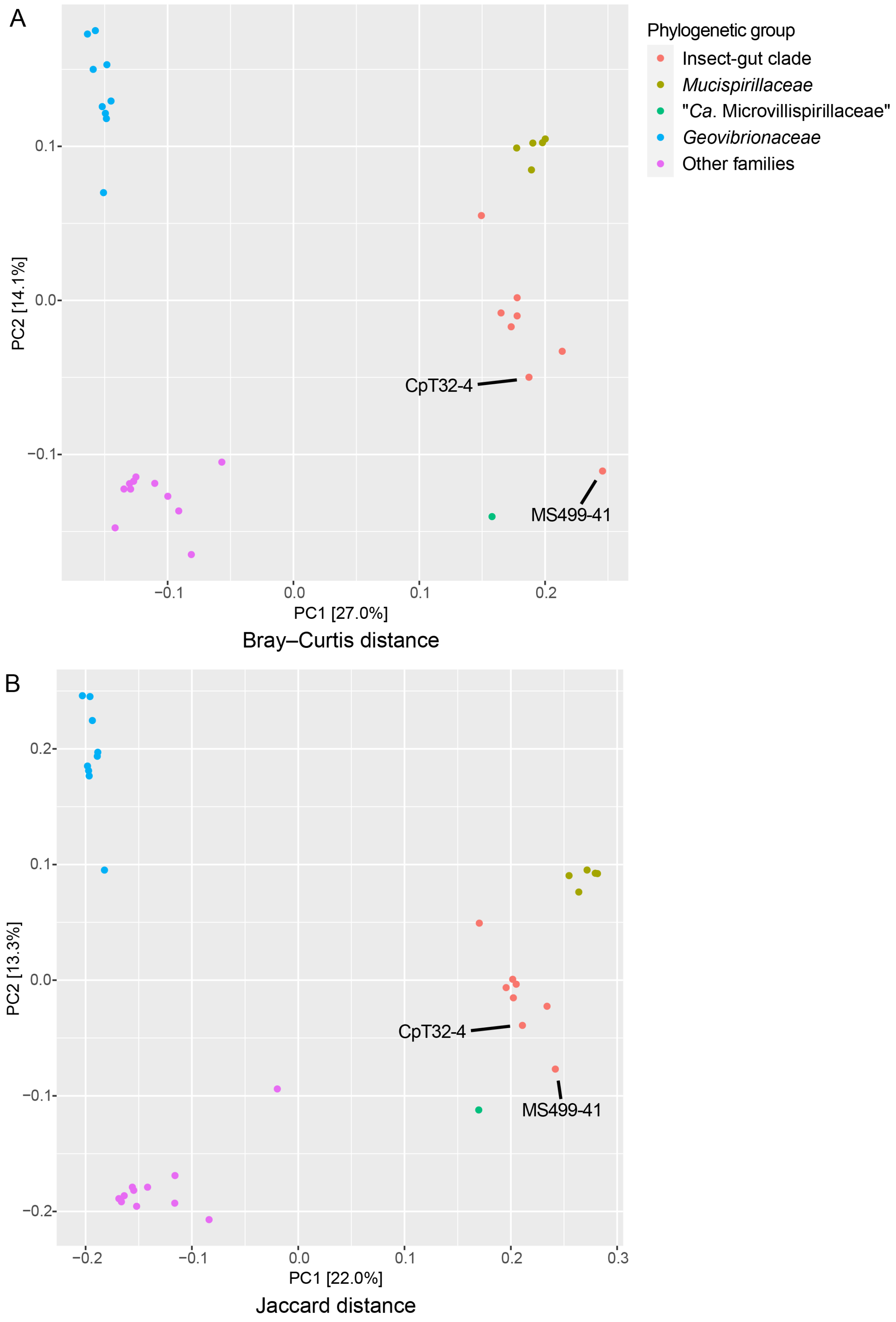


**Figure S10.** Similarity of genomic content among *Deferribacterota*. PCoA plots based on (A) Bray-Curtis distance and (B) Jaccard distance, calculated from orthogroup compositions. *Termitispirillum cryptocerci* (CpT32-4) and "*Candidatus* Termitispirillum reticulitermitis" (MS499-41) are indicated.

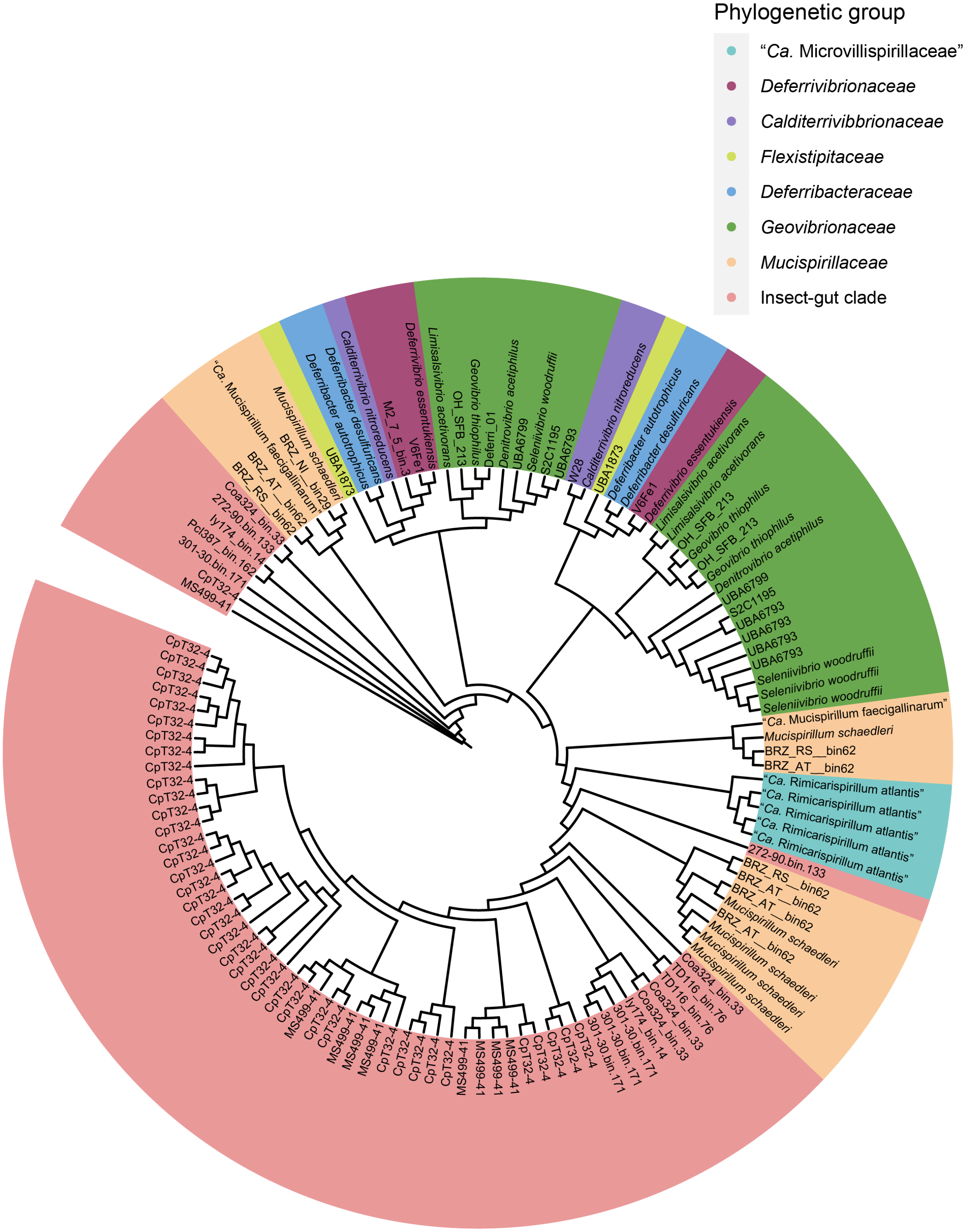


**Figure S11.** Phylogenetic diversity of the flagellin gene of *Deferribacterota*, including *Termitispirillum cryptocerci* (CpT32-4) and "*Candidatus* Termitispirillum reticulitermitis" (MS499-41). A maximum likelihood tree was constructed with the LG+F+G4 substitution model, based on 624 amino acid positions. Only genomes with the completeness >70% and contamination rate <5% were used as the sources of the flagellin genes.
